# 14-3-3 binding motif phosphorylation disrupts Hdac4 organized condensates to stimulate cardiac reprogramming

**DOI:** 10.1101/2023.11.20.567913

**Authors:** Liu Liu, Ienglam Lei, Shuo Tian, Wenbin Gao, Yijing Guo, Zhaokai Li, Ziad Sabry, Paul Tang, Y. Eugene Chen, Zhong Wang

## Abstract

Cell fate conversion is associated with extensive epigenetic and post translational modifications (PTMs) and architectural changes of sub-organelles and organelles, yet how these events are interconnected remains unknown. We report here the identification of a phosphorylation code in 14-3-3 binding motifs (PC14-3-3) that greatly stimulates induced cardiomyocyte (iCM) formation from fibroblasts. PC14-3-3 was identified in pivotal functional proteins for iCM reprogramming, including transcription factors and epigenetic factors. Akt1 kinase and PP2A phosphatase were a key writer and eraser of the PC14-3-3 code, respectively. PC14-3-3 activation induces iCM formation with the presence of only Tbx5. In contrast, PC14-3-3 inhibition by mutagenesis or inhibitor-mediated code removal abolished reprogramming. We discovered that key PC14-3-3 embedded factors, such as Hdac4, Mef2c, Nrip1, and Foxo1, formed Hdac4 organized inhibitory nuclear condensates. Notably, PC14-3-3 activation disrupted Hdac4 condensates to promote cardiac gene expression. Our study suggests that sub-organelle dynamics regulated by a post-translational modification code could be a general mechanism for stimulating cell reprogramming and organ regeneration.

**Highlights:**

1. A PC14-3-3 (phosphorylation code in 14-3-3 binding motifs) is identified in pivotal functional proteins, such as HDAC4 and Mef2c, that stimulates iCM formation.
2. Akt1 kinase and PP2A phosphatase are a key writer and a key eraser of the PC14-3-3 code, respectively, and PC14-3-3 code activation can replace Mef2c and Gata4 in cardiac reprogramming.
3. PC14-3-3 activation disrupts Hdac4 organized condensates which results in releasing multiple 14-3-3 motif embedded proteins from the condensates to stimulate cardiac reprogramming.
4. Sub-organelle dynamics and function regulated by a post-translational modification code could be a general mechanism in stimulating cell reprogramming and organ regeneration.

**Graphic abstract:** 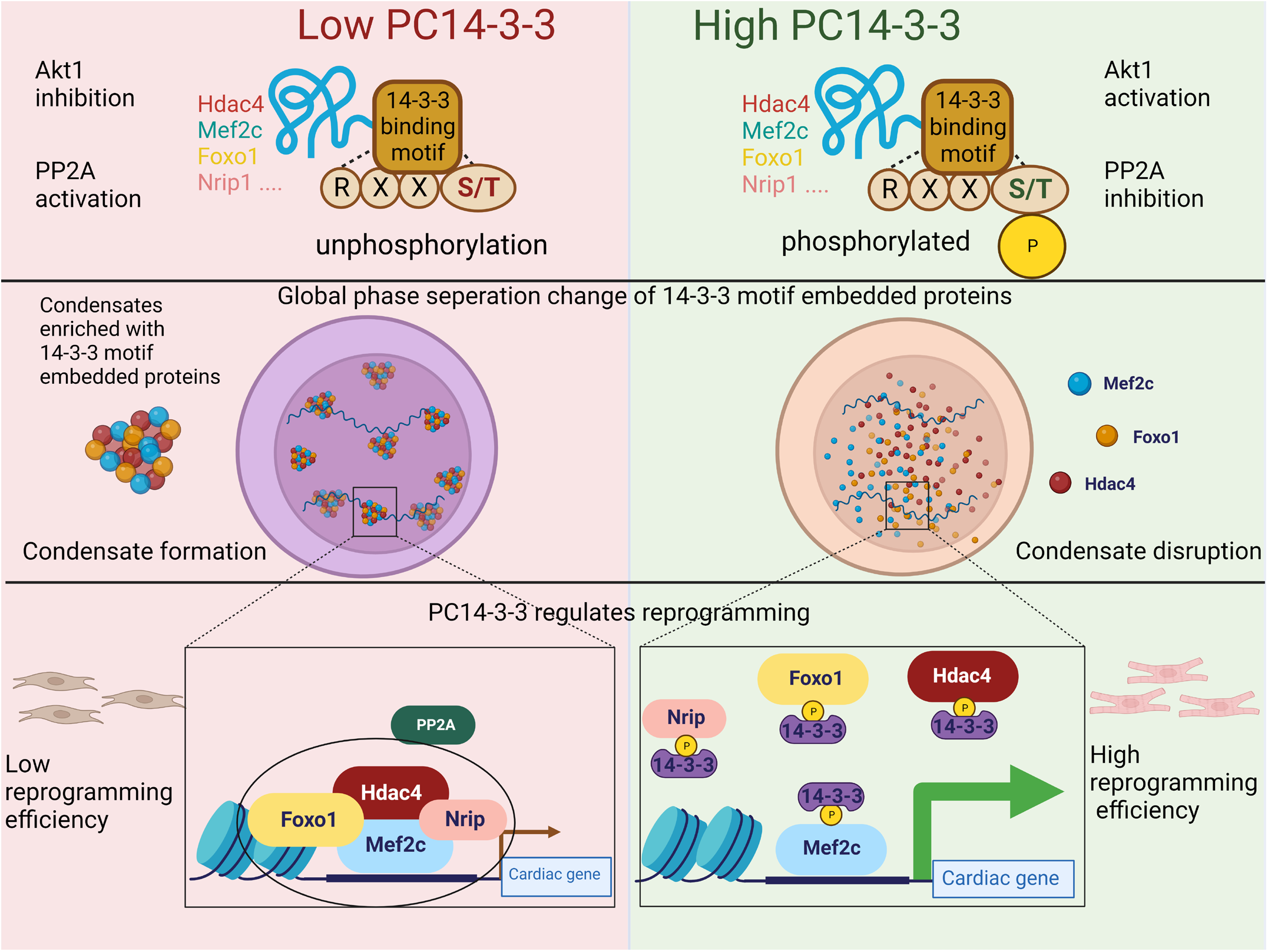

## Introduction

Heart disease is the leading cause of death in the United States and around the world. In heart injuries, such as myocardial infarction, millions of cardiomyocytes undergo irreversible necrosis and fibrosis. For patients who survive the myocardial infarction, the injured muscle tissues in the infarcted heart area are replaced with nonfunctional fibrotic scar tissues, which leads to severely weakened pumping function of the heart. There is a critical need for developing effective therapeutic strategies to preserve the pumping function after myocardial infarction. Direct reprogramming of fibroblasts into cardiomyocytes (CM)-like cells (called induced cardiomyocytes, iCMs) by introducing three cardiac transcription factors Gata4, Mef2C and Tbx5 (G, M and T) has emerged as an attractive strategy to repair damaged hearts (*1–3*). These iCMs, when generated in situ in an infarcted heart, integrate electrically and mechanically with surrounding myocardium, leading to a reduction in scar size and an improvement in heart function. However, low conversion rate, poor purity, and lack of precise conversion of iCMs remain significant challenges. Limited understanding of the molecular mechanisms of iCM reprogramming is a key obstacle preventing its effective clinical applications.

Cardiac reprogramming is a cell fate conversion process involving extensive epigenetic changes and post translational modifications (PTMs) (*4–9*). PTMs of histones have been extensively studied as a coding system to guide alterations in cell plasticity and cell fate, such as differentiation/programming and de-differentiation/reprogramming. Coordinated PTMs on chromatin function as a histone code and this code is added, read, and erased by epigenetic factors (*10*). In parallel with the PTMs coding system of histones, PTMs of free proteins are also essential for organ development and cell differentiation. Due to the complexity of PTMs on free proteins, most studies focus on specific PTM sites in a particular protein and generally define those PTMs as a feature belonging to the individual protein. However, the functional output of similar PTM on a broad range of motif embedded proteins as a coding system has not been explored. We propose that the histone code can be logically extended to include transcription factors and epigenetic factors termed as “PTM code”. This PTM code will be a complementary coding system to the genetic code to guide cell fate conversion, such as iCM reprogramming.

Cell fate conversion is also associated with extensive architectural changes of sub-organelles and organelles, yet how these organelle/sub-organelle dynamics are regulated and contributed to cell fate change is largely unknown. Biomolecular condensates (phase separation of proteins) are non-membrane bounded sub-organelles composed of heterogeneous or homogeneous proteins (*11, 12*). Recent studies indicate that epigenetic and transcription factors can aggregate into nuclear condensates and synergistically regulate gene expression (*13–15*). These condensates could play an essential role in cell fate change. Condensate formation typically requires phase separation region within the intrinsically disordered regions (IDRs) that don’t form a fixed three-dimensional structure (*16*). Notably, phosphorylation can alter the overall charge of IDR in the protein or alter the protein-protein interaction to regulate the condensate formation (*17*).

In this study, we report our identification of a phosphorylation code in 14-3-3 binding motif proteins (PC14-3-3) that guide cardiac reprogramming. Our initial screen has identified the presence of PC14-3-3 in pivotal functional proteins that stimulate iCM formation. We next show that PC14-3-3 code removal abolishes cardiac reprogramming. Furthermore, we also identified that Akt1 kinase and PP2A phosphatase were a key writer and a key eraser of the PC14-3-3 code, respectively. Activation of PC14-3-3 code by Akt1 expression and PP2A inhibition greatly enhanced reprogramming, inducing iCM formation with the presence of only Tbx5. Mechanistically, PC14-3-3 activation disrupts nuclear inhibitory Hdac4 condensates that trap other PC14-3-3 embedded factors which in turn releases transcription factors to promote cardiac gene expression. Our research reveals PC14-3-3 as an essential phosphorylation code regulates sub-organelle dynamics to stimulate cardiac reprogramming, which may have general significance in cell fate determination and cell fate change.

## Results

### Phosphorylation in 14-3-3 binding motifs is a phosphorylation code embedded in diverse classes of proteins that regulates cardiac reprogramming

To explore the potentially key role of phosphorylation in iCM induction, we first constructed a phosphorylation deficient mutant library for cardiac reprogramming factors. We picked the three primary factors MGT (Mef2c, Gata4 and Tbx5) as well as three other transcription factors (Hand2, Mesp1, and Nkx2-5) that play important functions in cardiac reprogramming. We attempted to include all the possible phosphorylation sites on those transcription factors, which included both the known (*18, 19*) and predicted ones (Table S1). Three softwares PhosphoSVM (*20*), NetPhos (*21*) and Musite (*22*) for phosphorylation prediction were used to identify 132 sites on all the transcription factors (TFs) with a high confident rate (Table S1). Therefore 132 known or predicted phosphorylation sites on these proteins were mutated into alanine to mimic dephosphorylated states.

Primary factors MGT or MGT plus either Hand2, Mesp1, or Nkx2-5 were introduced into MEF to induce cardiac reprogramming. TFs with dephosphorylation mimicking mutations were tested side by side with the wild type (WT) counterparts for their ability to affect iCM induction following the established iCM evaluation protocol (Fig. 1A). 14 dephosphorylation mimicking modification sites were observed to affect reprogramming based on the expression of Myh6 (Fig. 1B). Key mutants that dramatically affect reprogramming were also validated by measuring the cardiomyocyte structure gene cTnt protein expression with immunocytochemistry (ICC) (Fig. 1C&D) & (Fig. S1A). These results indicate that phosphorylation modifications of TFs are critical for cardiac reprogramming.

**Fig. 1.**
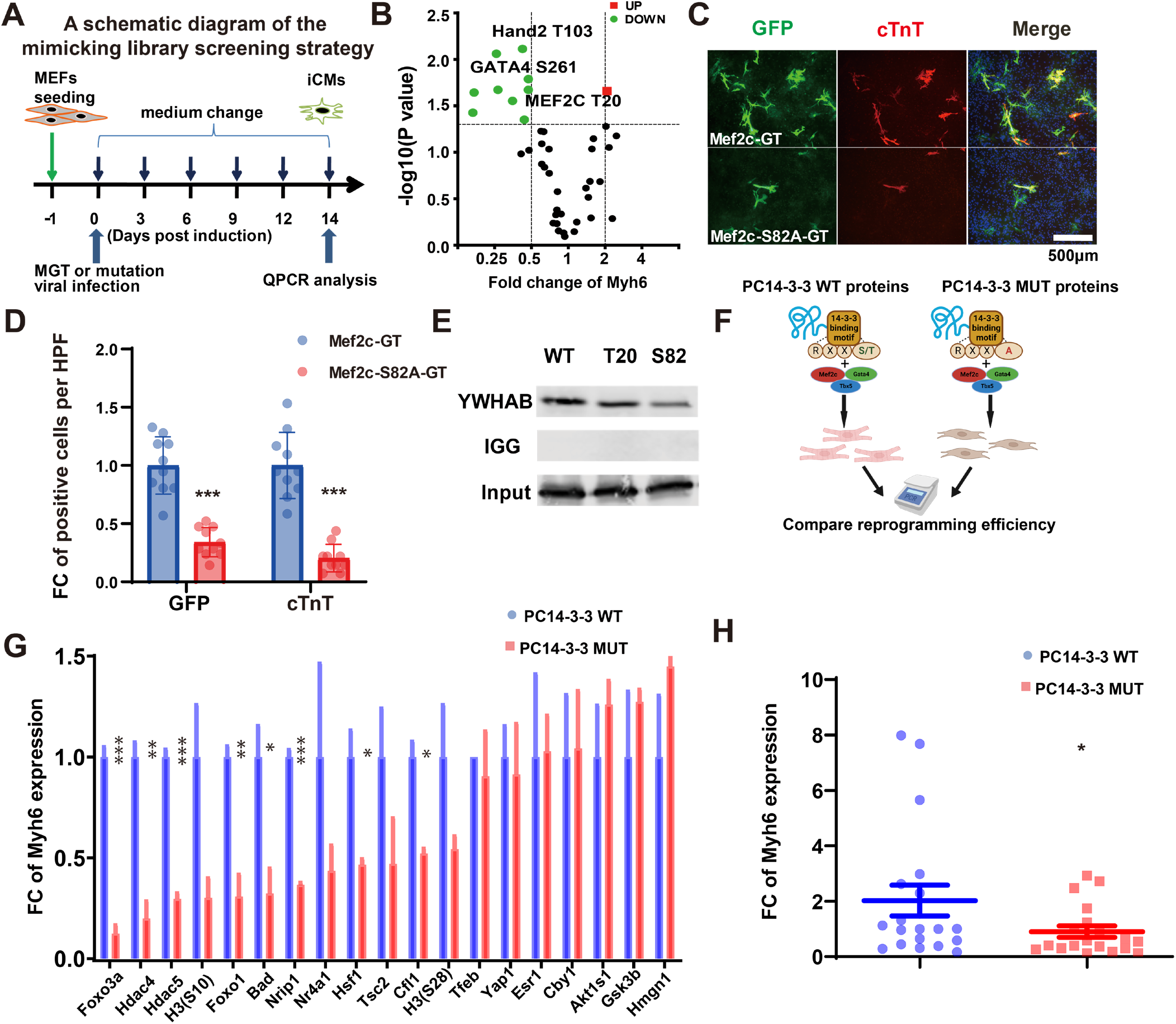
Phosphorylation in 14-3-3 binding motifs is a phosphorylation code embedded in diverse classes of proteins that can regulates cardiac reprogramming. (A) A schematic diagram of the phosphorylation screen strategy. Mutated individual gene was transduced into MEFs derived from a transgenic αMHC-GFP reporter mouse with retroviruses expressing. Medium was changed every 2 days. Reprogramming efficiency was evaluated 14 days after transduction by QPCR. (B) Myh6 expression measured after MEF induction with dephosphorylation mutation library. (C)&(D) Immunocytochemistry (ICC) of cardiac markers cTnt of S82A mutation construct or wt-transduced cells by fluorescence microscopy (100×). Mef2c-GT: Mef2c+Gata4+Tbx5. *P<0.05, **P<0.01, ***P<0.001, vs MGT. (E) Mutations of 14-3-3 binding motif phosphorylation disrupted 14-3-3 binding to Mef2c. HEK293 were transfected with 14-3-3 together with either wt or S82A or ST20A mutation of Mef2c. Mef2c was immunoprecipitated with anti-Mef2c antibody followed by western blotting and detection of 14-3-3 isoform ywhab-flag by flag antibody. (F) A schematic diagram of the 14-3-3 binding motif dephosphorylation mimicking screen strategy. Mutated individual gene was transduced into MEFs derived from a transgenic αMHC-GFP reporter mouse with retroviruses expressing MGT. Reprogramming efficiency was evaluated 14 days after transduction by QPCR as shown in A. (G) Relative expression of Myh6 in iCM reprogramming with WT and mutant PC14-3-3-containing proteins 14 days after MGT transduction. **P<0.01, vs WT. (H) Relative expression of Myh6 in iCM reprogramming with WT and mutant PC14-3-3-containing proteins 14 days after MGT transduction. Data are normalized to MGT+empty plasmids group. *P<0.05, **P<0.01, ***P<0.001, vs WT. FC : fold change

Next, we examined how these phosphorylation sites affected reprogramming. Since phosphorylation is well known for regulating the adjacent motif function, we mapped the phosphorylation site onto predicted motifs using the eukaryotic linear motif (ELM) resource (*23*). Intriguingly, the 14-3-3 binding motif was identified as the possible dominant motif in these TFs to regulate reprogramming (Fig. S1B). We further validated that Mef2c S82A mutant diminishes 14-3-3 interaction with the mutant as shown by our immunoprecipitation (IP) assays (Fig. 1E). Consequently, at least, the phosphorylation of 14-3-3 binding motifs within Mef2c could affect reprogramming.

We then examined the pathways related to the 14-3-3 binding motif-containing proteins during cardiac reprogramming with gene ontology (GO) analysis. Our analysis indicated that 14-3-3 binding motif-containing proteins were highly related to multiple pathways involved in cardiac reprogramming (Fig. S1C). Many transcription factors, epigenetic factors, and histone proteins related to heart development, muscle cell differentiation, and muscle cell apoptosis were also identified. To determine if phosphorylation of 14-3-3 binding motifs embedded in heart development or muscle cell differentiation proteins could contribute to regulate cardiac reprogramming, we next selected 19 key transcription factors, epigenetic factors and histone proteins carrying known 14-3-3 binding motifs to validate the possible function of their 14-3-3 binding motifs in cardiac reprogramming (Table S2). We created 14-3-3 binding motif dephosphorylation mimicking mutant protein as a control for each WT protein (Fig. 1F). Primary factors MGT plus WT or 14-3-3 binding motif mutant protein were introduced into MEF to induce cardiac reprogramming. Mutants were tested side by side with the WT counterparts for their ability to affect iCM induction following the established iCM evaluation protocol (Fig. 1A). 7 out of 18 tested protein mutants including those of Hdac4, Foxo1, Nrip1 and Bad, significantly inhibited cardiac reprogramming based on Myh6 and Actc1 expression (Fig. 1G & Fig. S1D). More importantly, compared to the WT protein, a general decrease in cardiac reprogramming was observed from the mutants (Fig. 1H & Fig. S1E). These observations indicate that phosphorylation of 14-3-3 binding motifs in these factors is the key phosphorylation event for cardiac reprogramming.

### 14-3-3 binding motif phosphorylation is required for and positively associated with cell fate conversion from fibroblasts to iCMs

Next, we determined the changes of 14-3-3 binding motif phosphorylation during reprogramming. We established an inducible reprogramming cell lines use cardiac-specific gene Myh6 promoter-driven fluorescence gene GFP as a marker to evaluate cardiac reprogramming (Fig. 2A). The polycistronic MGT and the transactivator to control of a tetracycline responsive promoter was introduced into this inducible cell line. Successfully reprogrammed Myh6-GFP+ cells can be separated from the unsuccessfully reprogrammed Myh6-GFP-cells by flow cytometry sorting (Fig. 2B). Since 14-3-3 binding motifs are broadly present in many proteins, we first examined the phosphorylation levels of all the 14-3-3 binding motifs during reprogramming using the phosphorylation-14-3-3 motif antibody (*24*). Indeed, the overall 14-3-3 binding motif phosphorylation is much higher in successfully reprogrammed CMs than non-reprogrammed cells (Fig. S2A). In order to study 14-3-3 binding motif phosphorylation dynamics during the reprogramming process, proteins from MGT and control vector infected MEF cells were harvested every two days up to day 14. The overall 14-3-3 binding motif phosphorylation was increasing during reprogramming in MGT infected cells especially from Day 4 to Day 12 (Fig. 2C). To determine whether a high level of phosphorylation in 14-3-3 binding motifs is unique to CMs among heart cells, neonatal CMs and CFs were also purified and the phosphorylation of 14-3-3 binding motifs was compared (Fig. S2B). Much higher phosphorylation in 14-3-3 binding motifs was observed in CMs than in CFs.

**Fig. 2.**
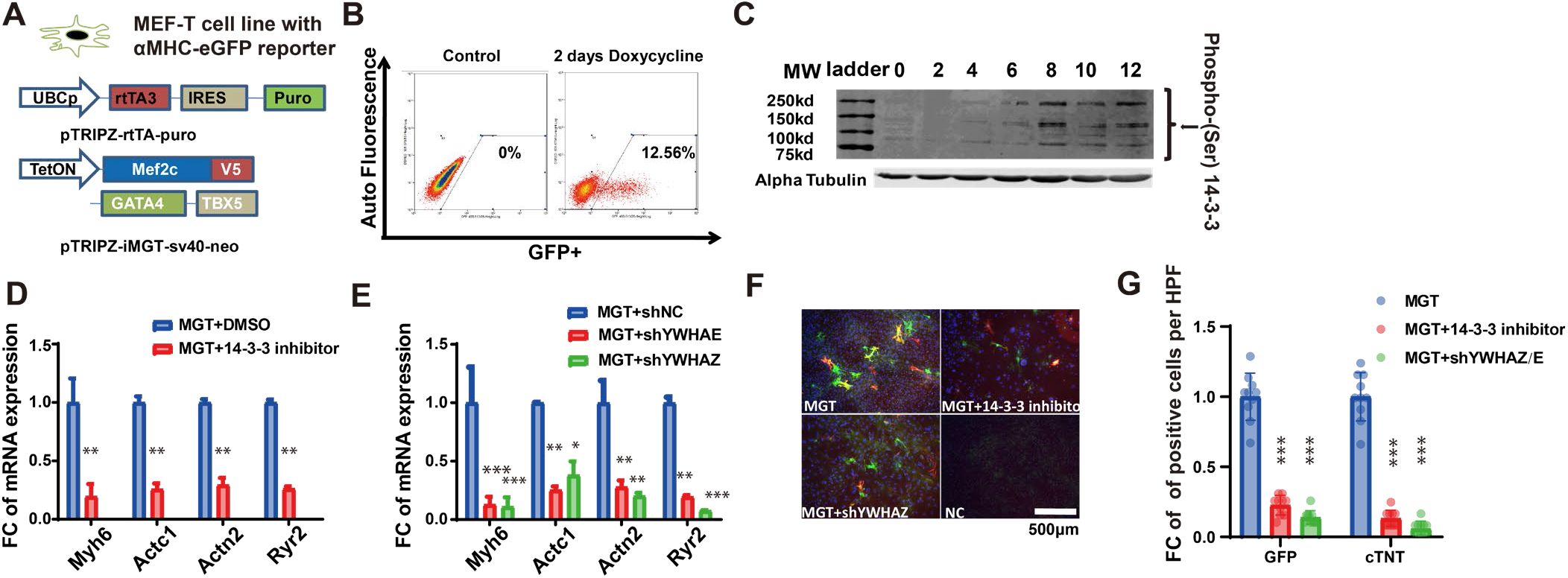
Phosphorylation in 14-3-3 binding motifs is required and positively associated with conversion from fibroblasts to iCMs. (A) A schematic diagram of establishing an inducible cardiac reprogramming cell line with cardiac-specific gene Myh6 promoter-driven GFP marker. (B) Successful reprogrammed Myh6-GFP+ cells and the unsuccessful reprogrammed Myh6-GFP-cells can be separated by flow cytometry sorting. (C) Detection of 14-3-3 binding motif phosphorylation by Western blot of protein lysates from MGT and control vector infection. MEF cells were harvested every two days until day 14. (D) 14-3-3 inhibitor (14-3-3 Antagonist I, 2-5) and (E) direct shRNA knockdown of 14-3-3 proteins significantly decreased reprogramming efficiency by QPCR. (F&G) Immunocytochemistry (ICC) and Quantification of cardiac markers of cardiac markers cTnT of MGT-transduced cells with 14-3-3 inhibitor or shRNA knockdown of 14-3-3 proteins by fluorescence microscopy. *P<0.05, **P<0.01, ***P<0.001,vs MGT.

A major function of the phosphorylated 14-3-3 binding motifs is to engage the interaction of the motif-containing proteins with its reader: 14-3-3(*25*). To further determine the role of phosphorylated 14-3-3 binding motifs in iCM formation, we applied 14-3-3 inhibitor (14-3-3 Antagonist I, 2-5) (Fig. 2D) and shRNA against 14-3-3 proteins YWHAE and YWHAZ (Fig. 2E & Fig. S2C) to disrupt the interaction of PC14-3-3 with 14-3-3s. Both 14-3-3 inhibitor and shRNAs against 14-3-3 significantly decrease reprogramming efficiency (Fig. 2F&G), indicating that the interaction of PC14-3-3-containing proteins with its reader 14-3-3 is essential for cardiac reprogramming. We also further validated the function of 14-3-3 in reprogramming by expressing a dominant negative K49E mutation in YWHAE (*26, 27*) during the reprogramming process. The results show that disrupting 14-3-3 function by K49E mutation also reduced cardiac reprogramming efficiency in the reprogramming cell line (Fig. S2D). In contrast, overexpression of 14-3-3 proteins did not induce much change in cardiac reprogramming efficiency (Fig. S2E). These results indicate that the phosphorylation in 14-3-3 binding motifs but not 14-3-3 protein themselves is key to cardiac reprogramming. Based on these results, we hypothesize that the phosphorylation of 14-3-3 binding motifs in many important functional proteins could be coordinately stimulated during cardiac reprogramming and this phosphorylation stimulation is essential for cardiac reprogramming. We define the phosphorylation acquired on those 14-3-3 binding motifs as a phosphorylation code of 14-3-3 binding motifs (PC14-3-3).

### PP2A inhibitor okadaic acid (OA) and Akt1 treatment leads to PC14-3-3 activation and drastically enhances reprogramming

The identification of the PC14-3-3 code prompted us to identify the key kinases (writers) and phosphatases (erasers) of the code to regulate reprogramming. Intriguingly, 6 out of those 8 14-3-3 binding motif embedded proteins identified in Fig. 1G that affected reprogramming are regulated by OA or PP2A (*28–37*), and 3 out of the 8 have been reported to be regulated by Akt1 (Table S3) (*31, 38–41*). Moreover, 7 out of 8 are regulated by either PP2A or Akt1. Therefore, we focused on PP2A and Akt1 as potential key eraser and writer, respectively.

Indeed, the addition of pharmaceutical PP2A inhibitor okadaic acid (OA) significantly activated PC14-3-3 code (Fig., 3A). To test whether there is a correlation between PC14-3-3 stimulation and reprogramming efficiency, we examined the effect of OA on reprogramming. OA treatment promoted higher cardiac troponin T (cTnT) and α-actinin expression and more sarcomere formation with up-regulation of cardiac genes (Fig. S3 B, C &D). PP2A inhibition also promoted higher spontaneous beating and calcium transient activities of iCMs derived from fibroblasts (Fig. S3 E&F). Consistent with these observations, PP2A knockdown by shRNA increased cardiac reprogramming efficiency (Fig. S3G&H).. Therefore, PP2A inhibition enhanced reprogramming mainly by activating PC14-3-3.

We next focused on Akt1 as the kinase for PC14-3-3. We analyzed the available database and/or the motifs activation correlation with PC14-3-3 binding motif phosphorylation (Fig. S3I). Candidates are listed based on the available database and/or the activated motifs correlation with PC14-3-3 binding motif phosphorylation (*42*). The analysis suggested that 70%, 74%, 20% and 14% of the PC14-3-3 code were likely to be written by Akt1, PKA, AMPK and CAMK, respectively. And Akt1 had the highest enrichment in the database. More importantly, Akt1 has been reported to enhance reprogramming (*41*).Therefore, Akt1 very likely enhanced reprogramming mainly by activating PC14-3-3.

We then examined the combinatorial effect of OA and Akt1 on PC14-3-3 activation and reprogramming. OA and Akt1 combination induced much higher PC14-3-3 activation than any individual treatment (Fig. S3A) and higher PC14-3-3 activation also led to higher reprogramming efficiency. Particularly, the four gene expression profiles (sarcomeric, contractility, ion channel genes Actn2, Actc1, Ryr2, and Myh6) indicated that reprogramming efficiency increased dramatically with OA and Akt1 treatment compared to the standard MGT procedure represented by a 75 fold increase of Myh6 expression (Fig. 3B). The PC14-3-3 activation by OA and Akt1 for two weeks also promoted iCM maturation as indicated by immunostaining against cardiac Troponin T (cTnT), the specific components of the sarcomere in CMs (Fig. 3C&D). Moreover, spontaneously contracting cells were apparent 1 week after PC14-3-3 activation treatment (Fig. 3E). Approximately 60 beating loci per well were identified in the PC14-3-3 activated group 4 weeks after transduction, 10-fold more than in the MGT control group. Consistent with more beating cells, PC14-3-3 activated iCMs showed significantly higher spontaneous Ca2+ oscillations with various frequencies (Fig. 3F). More importantly, YWHAZ/E knockdown significantly decreased Akt1+OA induced reprogramming (Fig. 3B-F). All our data and analyses indicated that although there could be diverse kinases and phosphatases for PC14-3-3 code, most of the motifs having significant effect on reprogramming can be phosphorylated by Akt1 and dephosphorylated by PP2A. Consequently, PP2A inhibitor OA and Akt1 synergistically led to PC14-3-3 activation and drastically enhanced iCM reprogramming. In contrast, PP2A activator DT-061 and/or Akt1 inhibitor MK-2206 treatment downregulated PC14-3-3 (Fig. 3G) and reprogramming efficiency in inducible reprogramming cell line (Fig. 3H).

**Fig. 3.**
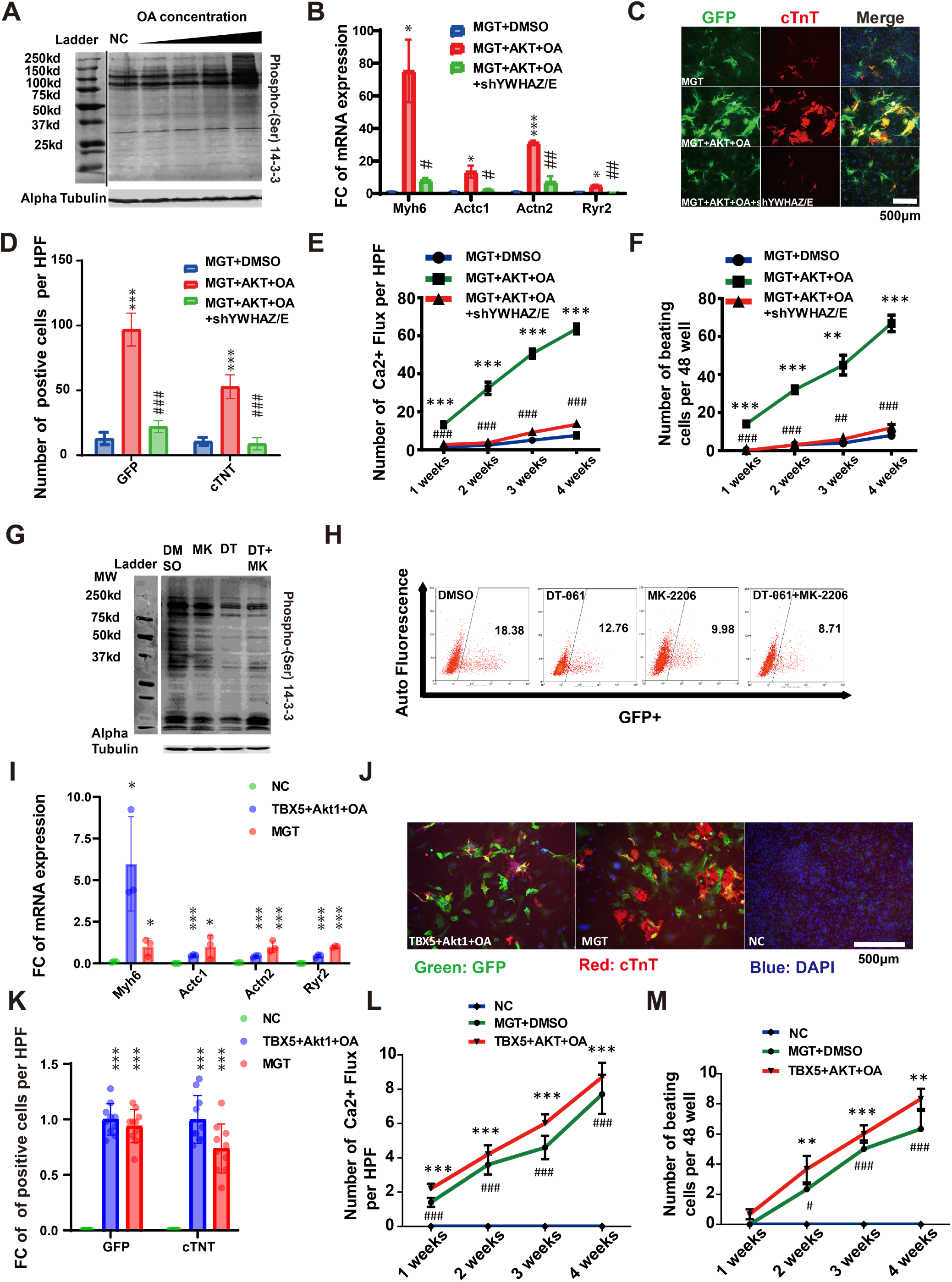
PP2A inhibitor okadaic acid and Akt1 treatment leads to PC14-3-3 activation and drastically enhances reprogramming. (A) Phosphorylation of 14-3-3 binding motifs was detected by Western blot of protein lysates from MEF cells with OA treatment. (B) Relative expression of cardiomyocyte (CM) marker genes of iCM reprogramming with Akt1 and OA treatment 14 days after MGT transduction with or without shRNA YWHAZ/E. Statistic analyses were performed between MEF with Akt1 and OA or DMSO-treated groups. *P<0.05,vs MGT. #P<0.05, ##P<0.01, vs MGT+Akt1+OA. (C&D) Immunocytochemistry (ICC) and Quantification of cardiac markers Tnt and Myh6-GFP of MGT-transduced cells with or without Akt1 and OA treatment with or without shRNA YWHAZ/E by fluorescence microscopy (100×). ***P<0.001, vs MGT. ###P<0.001,vs MGT+Akt1+OA. (E) Quantification of spontaneous Ca^2+^ oscillations cells per field with indicated viral infection treatment for 1 to 4 weeks (n=50 from 10 wells). ***P<0.001,vs MGT+DMSO, ###P<0.001,vs MGT+Akt1+OA. (F) Quantification of beating iCMs loci with indicated viral infection for 1-4 weeks (n=10). ***P<0.001,vs MGT+DMSO, ###P<0.001,vs MGT+Akt1+OA. (G) Phosphorylation of 14-3-3 binding motifs was detected by Western blot of protein lysates from MEF cells with or without MK-2206 (Akt1 inhibitor)+ DT-061 (PP2A activator) treatment. (H) Successful reprogrammed Myh6-GFP+ cells was detected by flow cytometry with or without MK-2206 (Akt1 inhibitor)+ DT-061 (PP2A activator) treatment in iCM cell line. (I) Relative expression of cardiomyocyte (CM) marker genes in iCM reprogramming with Akt1 and OA treatment 14 days treatment after Tbx5 transduction. *P<0.05, **P<0.01,***P<0.001, vs NC. (J&K) Immunocytochemistry (ICC) and Quantification of cardiac markers α-actinin of Tbx5 or control-transduced cells with or without Akt1 and OA treatment by fluorescence microscopy (400×). ***P<0.001, vs NC. (L) Quantification of spontaneous Ca^2+^ oscillations cells per field or (M) beating iCMs loci with Tbx5 or control-transduced cells with or without Akt1 and OA treatment for 1-4 weeks (n=10). *P<0.05, **P<0.01, ***P<0.001, #P<0.05, ##P<0.01, ###P<0.001,vs NC.

To further determine the key role of PC14-3-3 in cardiac reprogramming, we tested whether PC14-3-3 activation could replace the primary cardiac reprogramming factors Mef2c, Gata4, or Tbx5 and identified that only Tbx5 was required to achieve cardiac reprogramming with PC14-3-3 activation. Based on these results, we examined in detail the Akt1+OA+TBX5 vs MGT induced cardiac reprogramming. Under this single TF condition with Akt1+OA, a similar expression of functionally important cardiac genes Actn2, Actc1, Ryr2, and Myh6 was observed (Fig. 3I). Furthermore, immunostaining against cardiac Troponin T (cTnT), a specific component of the sarcomere in cardiomyocyte, clearly identified the sarcomeric structure (Fig. 3J&K), indicating that Akt1+OA+TBX5 was able to promote iCM maturation. OA+Akt1+Tbx5 treated cardiac fibroblasts also induced beating iCMs (Fig. 3L) with spontaneous Ca2+ oscillations (Fig. 3M). Together, these data indicated that PC14-3-3 renders two reprogramming factors Mef2c and Gata4 dispensable in cardiac reprogramming.

### PC14-3-3 activation disrupts Hdac4 organized condensates containing numerous 14-3-3 binding motif proteins

Our earlier study indicated that protein mutants including Hdac4, Foxo1, Nrip1 and Bad which depleted phosphorylation of 14-3-3 binding motifs significantly inhibited cardiac reprogramming (Fig 1). Among those proteins, the most effective regulator was Hdac4 (Fig. S1D). Compare to WT Hdac4, Hdac4 mutant with depleted phosphorylation of 14-3-3 binding motifs also largely abolishes reprogramming at protein levels (Fig. S4A,B&C), suggesting a key role of Hdac4 in PC14-3-3 mediated reprogramming. We therefore initiated the investigation of the molecular mechanism of PC14-3-3 mediated reprogramming by examining the function of Hdac4 and its PC14-3-3 motifs.

To determine how HDAC4 is regulated by PC14-3-3, we first examined the dynamic of Class IIa HDACs (Hdac4+5+9) proteins during cardiac reprogramming. Surprisingly, under normal iCM inducing conditions, endogenous Class IIa HDACs forms condensates in the nucleus (Fig. 4A&B&Fig. S4D). More importantly, PC14-3-3 activation by Akt1+OA decreased Class IIa HDACs condensates number and size, whereas PC14-3-3 repression by MK-2206 + DT-061 increased Class IIa HDACs condensates number and size. To determine directly whether HDAC4 condensate formation was regulated by PC14-3-3, we applied an optoIDR assay to further test PC14-3-3 function in Hdac4 condensate formation (*43*). The photoinducible, self-associating Cry2 protein was labeled with mCherry and fused to the IDR of WT Hdac4 or mutant Hdac4(Fig. 4C) which contain the 14-3-3 binding motifs (Fig. 4D). The fusion of a portion of the 14-3-3 binding motif mutant HDAC4-3SA-IDR to Cry2-mCherry facilitated the rapid formation of micron-sized spherical droplets upon blue light stimulation (Fig. 4E). However, WT HDAC4-IDR-Cry2-mCherry did not form condensates with blue light stimulation within 10 mins (Fig. 4E). Moreover, deletion of 14-3-3 binding motif and adjacent region IDR caused loss of Hdac4 condensate formation, especially those containing the deletion of adjacent IDR around the 14-3-3 binding motif phosphorylation sites S246 and S632 (Fig. S4E). These data indicate that HDAC4 does indeed form condensates and that the Hdac4 condensates are regulated by PC14-3-3.

**Fig. 4.**
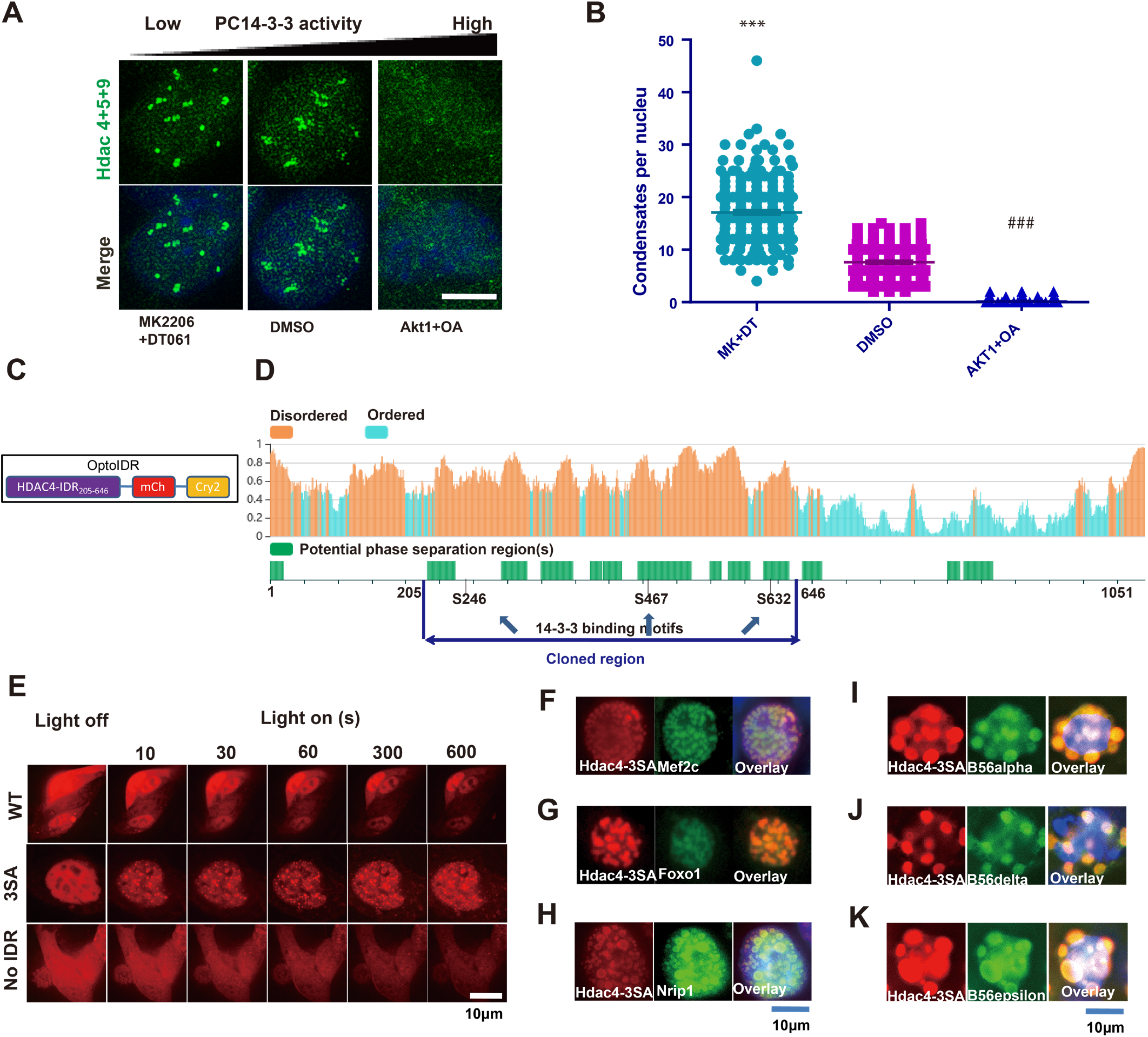
PC14-3-3 activation regulates Hdac4 organized condensates with multiple 14-3-3 motif embedded proteins. (A&B) Immunocytochemistry of endogenous Hdac4 in reprogramming cells treated with OA + Akt1 or MK-2206 (Akt1 inhibitor)+ DT-061 (PP2A activator) by fluorescence microscopy. Magnification image showing the Hdac4 condensates (the green dots) in a single reprogramming cell. (C&D) A schematic diagram showing Hdac4 14-3-3 binding motifs, intrinsically disordered regions (IDRs), potential phase separation regions, and the construction strategy for optoIDR assay. (E) Time-lapse images of the HE293T cell expressing Hdac4_205-646aa_-optoIDR with laser excitation. A droplet fusion event occurs in the Hdac4-3SA_205-646aa_ optoIDR group but not Hdac4_205-646aa_optoIDR. (F-H) Co-overexpression of Hdac4-3SA with PC14-3-3 embedded factors Mef2c (F), Foxo1 (G) and Nrip (H) showing that Hdac4 co-localized with these factors in the same condensate. (I-K) Co-overexpression of Hdac4-3SA with PP2A subunits showing that Hdac4 co-localizes with PP2A subunit B56alpha (I), B56delta (J) and B56epsilon (K) in the same condensate.

We next investigated the potential components within Hdac4 condensates. GO analysis of 14-3-3 binding protein showed that there is a cluster of proteins that are related to the nuclear body or granule (Fig. S4F). Intriguingly, optoIDR assay indicated that not only Hdac4, but also other PC14-3-3 embedded factors (Foxo1 and Mef2c) can regulate both reprogramming and condensate formation through PC14-3-3 activation (Fig. S4G-J). Most importantly, co-overexpression of PC14-3-3 mutant versions of Hdac4 (Hdac4-3SA) with PC14-3-3 embedded proteins in HEK293 cells show that Mef2c, Foxo1, or Nrip1 could co-localize with Hdac4 in the same puncta (Fig. 4F-H). We further screened PP2A subunits and observed that PP2A subunits B56alpha, B56delta and B56epsilon co-localized within Hdac4 condensates (Fig.Fig. 4I-K), suggesting that Hdac4 organized condensates are heavily regulated by PC14-3-3. These results imply that PC14-3-3 activation serves as a general mechanism to regulate the formation and dynamics of 14-3-3 motif embedded protein condensates, which are organized by Hdac4 and contains Mef2c, Foxo1 and Nrip1.

### Disruption of Hdac4 condensates is responsible for PC14-3-3 stimulated cardiac reprogramming

To determine if HDAC4 condensate disruption is responsible for PC14-3-3 stimulated cardiac reprogramming, we first generated several 14-3-3 permanent binding forms of Hdac4 to replace the 3SA mutants with R18 (Fig. 5A & Fig. S5A). R18 is an unphosphorylated peptide that binds to 14-3-3 proteins with high affinity (44, 45). We reason that if the effect of HDAC4 3SA mutant on inhibiting cardiac reprogramming is due to the disrupted binding of Hdac4 to 14-3-3, replacing the mutant 14-3-3 binding motifs with a peptide that binds strongly to 14-3-3 proteins will largely abolish the inhibitory effect. Therefore, the three 14-3-3 mutant binding motifs of Hdac4 3SA were replaced with an R18 peptide individually or collectively. The combined three binding motif mutate construct Hdac4-3R18 were introduced into MEF cells with MGT. WT and PC14-3-3 mutant forms of Hdac4 in the MEF cells show an obvious difference in cellular distribution with immunostaining (Fig. 5B&C). Unlike WT Hdac4 which distributed evenly and had no condensates in the cytosol, mutant Hdac4 only existed in nucleus and formed obvious biomolecular condensate-like puncta in the nucleus. Drastic dissociation of Hdac4 condensates were observed with the over-expression of Hdac4-3R18 further suggesting that 14-3-3 binding motifs are involved in the regulation of Hdac4 condensates (Fig. 5B). Moreover, Hdac4-3R18 indeed abolishes the inhibitory effect of Hdac4-3R18 in cardiac reprogramming (Fig. 5D). Consistent with the fact that PC14-3-3 activation disrupted Hdac4 condensates, other proteins within the condensates also lost their localization with Hdac4 condensates, such as Mef2c (Fig. 5C & Fig. S5C). Taken together, these results indicate that HDAC4 condensate formation is directly regulated by PC14-3-3 and is a key target of PC14-3-3 to stimulate cardiac reprogramming.

**Fig. 5.**
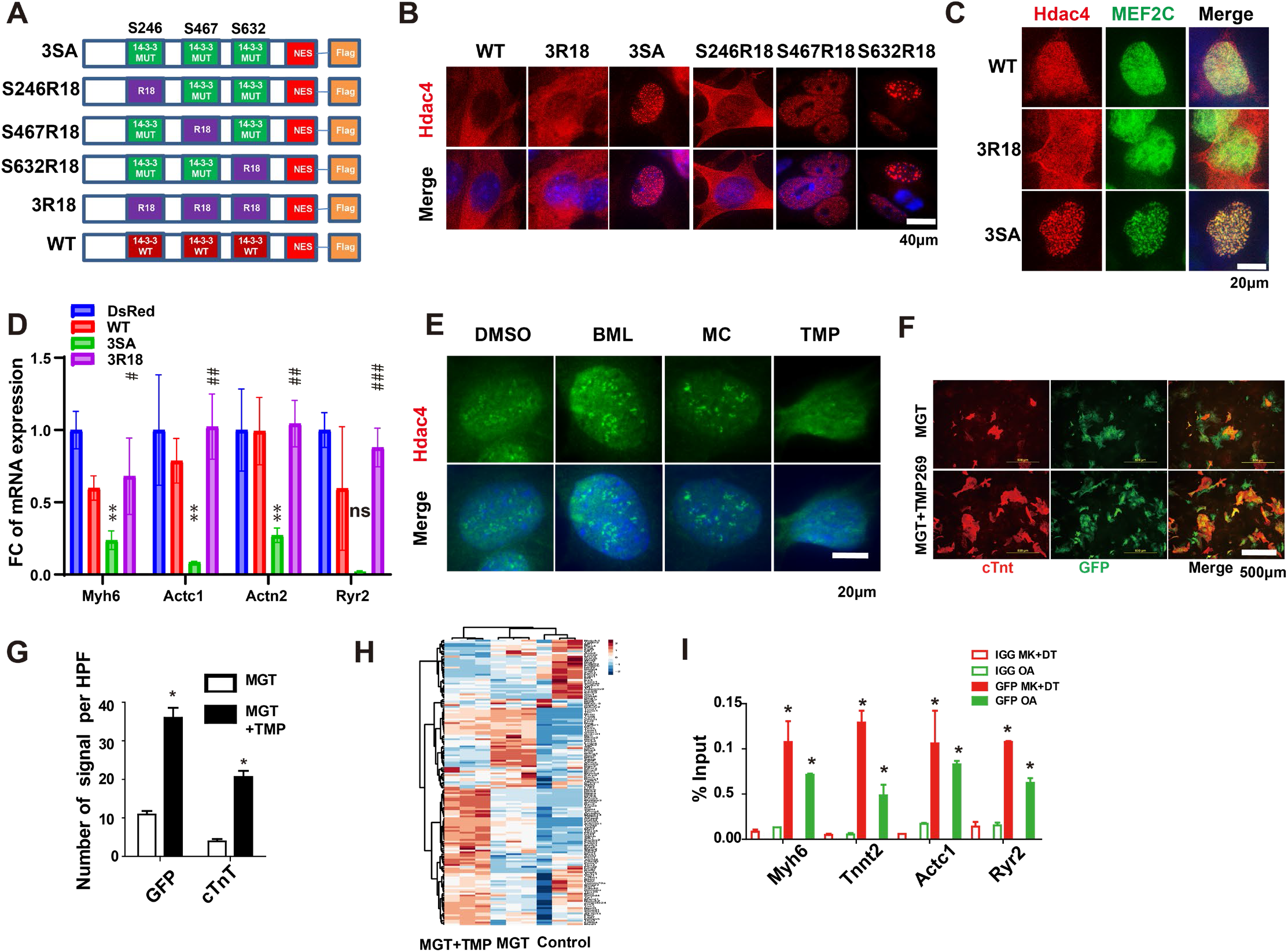
PC14-3-3 dissociates Hdac4 condensates to facilitate cardiac reprogramming. (A) A schematic diagram of Hdac4 mutants in 14-3-3 binding motifs combined with replacement of a permanent binding peptide R18. (B) Immunostaining of specific mutant Hdac4 proteins in reprogramming cells. (C) Co-overexpression of Hdac4 mutants with PC14-3-3 embedded factors Mef2c showing that R18 dissociates Hdac4 co-localized with MEF2C in the same condensate. (D) Relative expression of CM marker genes during iCM reprogramming with expression of Hdac4 mutants 3R18 and 3SA. *P<0.05, **P<0.01, ***P<0.001, WT vs 3SA. #P<0.05, ##P<0.01, ###P<0.001, 3SA vs 3R18. (E) Immunocytochemistry (ICC) of Hdac4 in iCM cells with Hdac4 overexpression and Hdac4 inhibitor treatment. Among Hdac4 inhibitors, TMP269, but not MC1568 and BML210, dissociates Hdac4 condensates. Immunocytochemistry (ICC) by fluorescence microscopy (F) and quantification (G) of cardiac markers cTnt and Myh6-GFP of MGT-transduced cells with Hdac4 inhibitor TMP269 treatment (400×). (H) Heat map showing the effect of Hdac4 inhibitor TMP269 on representative cardiac-related gene expression (GO: 0048738) among MGT-transduced cells. (I) ChIP-qPCR of Hdac4-GFP at cardiac gene loci from iCM cells treated with OA (PC14-3-3 activation) or MK2206 +DT-061 (MK+DT) (PC14-3-3 inhibition). n=3, *P<0.05, vs IGG.

It is also possible that PC14-3-3 stimulated cardiac reprogramming may be due to its effect on Hdac4 subcellular localization but not condensate regulation because phosphorylation of S246 can induce Hdac4 translocation to the cytosol (*46, 47*). To differentiate between these two possibilities, we constructed permanent nucleus localized Hdac4 forms by deleting the C-terminal nuclear export signal (NES)(*48*) on Hdac4 (Fig. S5A). The Hdac4-3R18-NES expression in nucleus did not lead to condensate formation in MEF (Fig. S5B). Moreover, Hdac4-3R18 expression indeed abolished the inhibitory effect of Hdac4 in cardiac reprogramming (Fig. S5D). This result reveals that the effect of PC14-3-3 on cardiac reprogramming is by regulating Hdac4 condensate dynamics but not Hdac4 subcellular localization. In addition, among the single 14-3-3 binding motif mutations of Hdac4, we observed significant changes in cardiac gene expression with Hdac4-1R18-246-2SA and Hdac4-1R18-632-2SA but not Hdac4-1R18-467-2SA, implying that PC14-3-3 on S246 and S632 is more important for cardiac reprogramming (Fig. S5E&F).

To independently determine the role of Hdac4 condensate dissociation in reprogramming, we screened and tested various Hdac4 inhibitors in condensate formation and cardiac reprogramming. Among the Hdac4 inhibitors tested, TMP269 but neither MC1568 nor BML210 dissociated Hdac4 condensates in iCM cells (Fig. 5E). In parallel, TMP269 enhanced cardiac reprogramming whereas MC 1568 and BML210 modestly inhibited it (Fig. S5G). HDAC4 inhibitor TMP269 improved MGT-mediated iCM formation by enhancing higher Myh6-GFP and cardiac troponin T (cTnT) expression (Fig. 5F&G). TMP269 also promoted higher calcium transient activities in iCMs derived from neonatal cardiac fibroblasts (Fig. S5H). Consistent with these observations, TMP269 treatment also promoted cardiac gene activation and down regulation of fibroblast genes at genome level (Fig. 5H & Fig. S5I). These results further support the conclusion that PC14-3-3 activation stimulates MGT-mediated iCM formation through HDAC4 condensate disruption.

We finally examined the function of Hdac4 condensates in regulating chromatin and cardiac gene expression. We examined if PC14-3-3 directly regulated Hdac4 at cardiac gene promoters during iCM formation. Cells overexpressing GFP-Hdac4-NES-deletion were used for ChIP assays in an iCM cell line using GFP antibodies (Fig. S5J). OA or MK2206+DT-061 were used to activate or repress PC14-3-3 activation. Hdac4 was localized to cardiac gene promoters including Myh6, Tnnt2, Actc1 and Myocd, and PC14-3-3 inactivation by MK2206+DT-061 enhanced Hdac4 binding (Fig. 5I). In contrast, PC14-3-3 activation with OA reduced Hdac4 binding. We also tested the H3K27ac on those promoters as a marker for gene activation and observed that H3K27ac was negatively associated with Hdac4 recruitment (Fig. S5K). These results show that Hdac4 binding to the cardiac promoter within the Hdac4 condensates induces a repressive effect for cardiac gene expression and PC14-3-3 activation disrupts the repressive effect.

Together, these data illustrate that without PC14-3-3 activation, Hdac4 inhibitory condensates trap other 14-3-3 binding motif carrying proteins inside the condensates and repress gene expression. In contrast, PC14-3-3 activation disrupts the repressive Hdac4 condensates and leads to the release of transcription/reprogramming factors, which may work synergistically to induce cardiac gene expression for cardiac reprogramming.

## Discussion

The key discovery of this research is the identification of the PC14-3-3 and its critical role in cardiac reprogramming. Inspired by the histone code concept, PC14-3-3 is defined as the coordinated phosphorylation modification at the 14-3-3 binding motifs within 14-4-3 binding proteins. The major characteristics of the PC14-3-3 code are: (1) the code is embedded within the 14-3-3 binding motifs in a large array of diverse proteins, which include transcription factors, epigenetic factors, and various pathway factors. The key factors carrying the PC14-3-3 that play a critical role in cardiac reprogramming are Hdac4, Mef2c, Foxo1,and Nrip; (2) the code is recognized and engaged by the reader 14-4-3 proteins; (3) the code is added or removed by multiple kinases and phosphatases with Akt1 and PP2A being the major identified writer and eraser in cardiac reprogramming; (4) a key function of the PC14-3-3 code is to induce a global phase change of 14-3-3 motif embedded proteins like Hdac4, Mef2c, Nrip1 and Foxo1 from a condensate phase to a free phase, that is, the activation of PC14-3-3 disrupts Hdac4 condensates and releases Mef2c, Foxo1 and Nrip1 for proper cardiac gene expression.

In this study, we identify Akt1 kinase and PP2A phosphatase as key writer and eraser of the PC14-3-3 code, respectively. Among the initial seven PC14-3-3-containing proteins affecting reprogramming, five are regulated by okadaic acid (OA) or PP2A and two are regulated by Akt, and 6/7 are regulated by either PP2A or Akt1. Moreover, PP2A inhibitor OA and Akt1 synergistically leads to PC14-3-3 activation, drastically enhance iCM reprogramming, and render Mef2c and Gata4 dispensable for reprogramming. In contrast, shRNA knockdown of 14-3-3**ε** (YWHAE) and 14-3-3**ξ** (YWHAZ) largely abolishes Akt1 and OA stimulated reprogramming. Furthermore, PP2A spatially co-localizes with PC14-3-3 regulated HDAC4 condensates. Nevertheless, aside from Akt1 and PP2A, we expect that there will be numerous writers and erases of the PC14-3-3 code, similar to those of histone code.

An important novel discovery is that PC14-3-3 very likely regulates cardiac reprogramming at a sub organelle level (condensates) via spatial protein interactions. Our investigation identifies the presence of Hdac4 condensates and its dynamic changes during cardiac reprogramming. PC14-3-3 activation decreases the size and number of Hdac4 condensates whereas PC14-3-3 inhibition increases it. Hdac4 14-3-3 binding motifs are localized in the adjacent condensate regulation IDR region. Mutations in the Hdac4 14-3-3 binding motifs greatly affects Hdac4 condensate dynamics. Our studies reveal that Hdac4 inhibits transcription and Hdac4 condensates are associated with transcriptionally inactive chromatin areas. Importantly, mutations in Hdac4 14-3-3 binding motifs show strong correlation between Hdac4 condensate formation and inhibition of cardiac reprogramming. Inducing PC14-3-3 by 14-3-3 permanent binding forms of Hdac4 also further shows strong correlation between inhibition of Hdac4 condensate formation and cardiac reprogramming. Finally, we have identified that among Hdac4 inhibitors, TMP269, but neither MC1568 nor BML210, dissociates Hdac4 condensates. In parallel, TMP269 enhances cardiac reprogramming whereas MC 1568 and BML210 inhibit it. These results lead us to conclude that Hdac4 condensate dissociation is responsible for PC14-3-3 stimulated reprogramming.

An interesting finding is that numerous factors, such as Mef2c, Nrip1 and Foxo1, co-localize within the Hdac4 condensates. This finding is consistent with the notion that Hdac4/5 can physically interact with Mef2c(*49, 50*), Nrip1(*51*) and Foxo1(*52, 53*). Also, PC14-3-3 appears in the adjacent condensate regulation IDR region of those factors. Our studies imply PC14-3-3 code embedded proteins may be involved in a distinct layer for chromatin and gene regulation. PC14-3-3 code embedded proteins spatially present in the same biomolecular condensates could provide rationality why seemingly discrete modifications or signals can have a unified function in cell fate change such as cardiac reprogramming. Our study indicates that PC14-3-3 activation induces a global phase change of 14-3-3 motif embedded proteins like Hdac4, Mef2c, Nrip1 and Foxo1, and by regulating the PC1-3-3 activation of different proteins, it likely induces a multilevel regulation of this condensate and its function. PC14-3-3 activation serves as a general mechanism to regulate the composition or composition activation in the condensates. It also worth noting the presence of numerous TF and regulators within the PC14-3-3 condensates. The disruption of Hdac4 condensates may not only diminish the Hdac4 activity, but also increases the activities of Mef2c, Nrip1, and Foxo1. Moreover, the presence of Mef2c in the condensates also explains at least partially why PC14-3-3 activation replaces Mef2c and Gata4 over-expression for cardiac reprogramming. Overall, the identification of the composition and dynamics of Hdac4 condensates during cardiac reprogramming and their regulation by PC14-3-3 has provided novel mechanistic insights into cardiac reprogramming.

This study implies that coding systems based on motifs and PTM modifications, such as PC14-3-3, very likely serve as a general mechanism in the biological system. We predict that a PTM code has the following characteristics: 1. the coding information is dependent on both motif sequences and PTM modifications. 2. This type of code possesses a collaborative function for a specific biological process by connecting the diversity of its embedded proteins. 3. Naturally, the code’s collaborative function is executed at multiple levels but very likely spatiotemporal connected. In this report, we identify that PC14-3-3 function depends on the phosphorylation of 14-3-3 binding motifs, PC14-3-3 activation play a key role in cardiac reprogramming, and its function is executed by disrupting Hdac4 condensates which connect Hdac4, numerous transcription factors, and possible chromatin. Our studies also show that PC14-3-3 activation also induces endothelial cell and smooth muscle cell trans-differentiation from fibroblasts (data not shown). These studies imply that PC14-3-3 activation represents a general mechanism in cell fate changes beyond CM reprogramming. We are optimistic that PTM code studies within free proteins will provide novel insights into biological processes, particularly cell fate establishment and cell fate change, which may reveal new targets for organ disease and regenerative medicine.

## Acknowledgements

This research was supported in part by R01HL139735 and R01HL163672 to ZW and HL109946 and HL159900 to YEC. Graphic abstract and Figure 1F were created with the help from BioRender.com. We thank Drs. Margaret V. Westfall, Jiandong Liu, Gregory Cartee, and Weidong Wang’s constructive suggestion to the manuscript.

## Author contributions

L.L., I.L and Z.W., conceptualization; L.L., and Z.W., data curation; L.L., I.L and Z.W., formal analysis; L.L., and Z.W., validation; L.L., I.L and Z.W., visualization; L.L., writing-original draft; L.L. and I.L, software; L.L., I.L and Z.W., methodology; L.L., I.L, S.T, W.G, Y.G, Z.L, Z.S, P.T, Y.C and Z.W., resources; L.L., I.L, S.T, W.G, Y.G, Z.L, Z.S, P.T, Y.C and Z.W.; L.L., Y.C and Z.W., project administration; L.L., I.L, S.T, W.G, Y.G, Z.L, Z.S, P.T, Y.C and Z.W., editing; L.L., Y.C and Z.W., supervision; Z.W., funding acquisition.

## Competing interests

The authors declare no competing interests.

## Materials and Methods

### Mouse lines

The α-MHC-GFP transgenic mice were used to derive MEFs and NCFs as described previously (*54–56*). αMHC-Cre/Rosa26A-Flox-Stop-Flox-GCaMP3 mice was purchased from Jackson Laboratories (011038, 014538). All animal related procedures were approved by the Institutional Animal Care and Use Committee of the University of Michigan and are consistent with the National Institutes of Health Guide for Use and Care of Animals.

### Plasmids

The pMXs based retroviral polycistronic vector encoding Mef2c, Gata4, Tbx5 were provided by Dr. Li Qian’s lab(*57*). Mutation constructions of cardiac genes Mef2c, Gata4, Tbx5, Hand2 and Nkx2-5 were constructed by overlap PCR strategy. The main primers used for the cloning was list in Supplementary Table S5. Most of the genes related with 14-3-3 were acquired from Addgene and cloned into pMX plasmids. The plasmids purchase from Addgene what listed in Supplementary Table S4. The 14-3-3 permanent bind forms of Hdac4 fragments of Hdac4-3R18 were synthesized by Genscript. The shRNA lentivirus vector for 14-3-3 isoforms, Ppp2cb were from Vector Core of University of Michigan.

### Inducible fibroblast cell line construction

This inducible fibroblast cell line construction was generated from α-MHC-GFP transgenic mice (*58*). All the Lentiviral Transfer Plasmids was revised from pTripz backbone. The polycistronic MGT construct under the control of a tetracycline responsive promoter for temporal control of MGT reprogramming factor expression and constitutive expression neomycin resistance gene (neo) was cloned. The second plasmid could express transactivator (reverse tetracycline-controlled transactivator) to induce tetracycline responsive promoter activation and constitutive expression puromycin resistance gene (puro) expression plasmids. The third plasmid was revised from retrovirus pMX to express Hdac4 and hygromycin resistance gene. Targeted plasmids was packaged into lentivirus or retrovirus, selected with G418 (750 µg/ml, invitrogen), puromycin (4ug/ml, invitrogen) or hygromycin B (1000 µg/ml, invitrogen) to generate MEF cell line (icMEF) that can be reprogrammed simply by the addition of doxycycline to the culture media.

### Inhibitor information

Okadaic Acid (10011490), TMP269 (17738), MC 1568 (16265), BML-210 (10005019), MK-2206 (11593) were purchased from Cayman. DT-061 (S8774) was purchased from SELLECK CHEMICALS. 14-3-3 Antagonist I, 2-5 (100081) was purchased from Sigma.

### Retrovirus and lentivirus preparation

75% confluent 10 cm dish of Platinum-E (Plat-E) Retroviral Packaging Cell Line (cell biolabs) were transfected with totally 10 μg retrovirus vectors using Lipofectamine 2000 (Thermo Fisher Scientific) in 1.5 ml Opti-MEM (Thermo Fisher Scientific). After 24 hours, medium was changed with 10 ml fresh DMEM medium with 10% FBS. Viral medium was collected twice 48 hours and 72 hours after transfection. The Viral medium was filtered through a 0.45-mmcellulose filter and was added 1/5 vol of 40% PEG8000 solution. The mixture was kept at 4 °C overnight and spun at 3000 g, 4 °C, 30 minutes to get concentrated. Virus was resuspended by fresh MEFs medium with 8 μg/ml polybrene (Sigma).

Similarly, 10 μg lentiviral vectors with 6 μg psPAX2 and 4 μg pMD2.G were packaged into an 60% confluent 10 cm plate of HEK293T cells (ATCC) with 0.75 ml Opti-MEM by using Lipofectamine 2000 (Invitrogen) with another 0.75 ml Opti-MEM with DNA mixture. 4-6 hours later, Opti-MEM was changed by 10 ml fresh MEFs medium. The virus was collected as descried before.

### Direct reprogramming of fibroblasts to iCMs

Preparation of MEFs (isolated at E13.5) was previously described (*55*). NCFs were isolated from P2-P3 α-MHC-GFP transgenic or αMHC-Cre/Rosa26A-Flox-Stop-Flox-GCaMP3 mice as reported previously (*54, 56*).

The optimized protocol of direct cardiac reprogramming was described obviously (*59*). Briefly, fresh fibroblasts were seeded on tissue culture dishes at a density of 10,000 cells/cm^2^ before virus infection. Fibroblasts were infected with fresh viral mixture containing 8 μg/ml polybrene (Sigma) 24 hours after seeding. Twenty-four hours later, the viral medium was replaced with induction medium composed of DMEM/199 (4:1) (Gibco, Thermo Fisher Scientific) containing 10% FBS, 1% penicillin/ streptomycin and 10 μl/ml GlutaMAX (Gibco, Thermo Fisher Scientific). Medium was changed every 2–3 days with or without indicated chemicals for two weeks before cells were examined. 1 μg/ml puromycin (SIGP8833-25MG, Sigma) was added into the medium 3 days after infection to eliminate cells without infection.

For spontaneous beating and calcium transient assessment experiments, induction medium was replaced every 2-3 days by mature medium containing StemPro-34 SF medium (Gibco, Thermo Fisher Scientific), GlutaMAX (10 μl/ml, Gibco, Thermo Fisher Scientific), ascorbic acid (50 μg/ml, Sigma), recombinant human VEGF165 (5 ng/ml, R&D Systems), recombinant human FGF10 (25 ng/ml, R&D Systems), and recombinant human FGF basic146 aa (10 ng/ml, R&D Systems) for another 2 weeks as previously described(*60*).

### Western blot

Proteins were extracted from cells by adding lysis buffer and centrifuged at 4 °C for 15 minutes at 12,000 rpm. The membranes were blocked with 4% BSA for 1 hour at room temperature and then incubated with primary antibodies over night at 4 °C. After 3 washings with TBST, the membranes were incubated with appropriate secondary antibody in TBST solution for another 1 hour at room temperature. After 3 washings, the membranes were scanned and quantified by Odyssey CLx Imaging System (LI-COR Biosciences, USA).

### Immunocytochemistry

Anti-HDAC4 + 5 + 9 antibody (Abcam, ab131524) is the antibody used to detect the endogenous condensate formation. This antibody directed target the 14-3-3 bind motif of Hdac4 around phosphorylated site S246. Hdac4-flag antibody (Sigma F1804) and H3K27ac (Abcam ab245911) or Pol2 Ser5 (Abclonal AP0828) antibodies V5 (Abcam ab9116). Mef2c (CST 5030), For Foxo1-HA and Nrip1-HA (ab9110). cTnT (Thermo Fisher Scientific), and GFP (Thermo Fisher Scientific) Cells were firstly fixed with 4% formaldehyde for 15 minutes. Then they were permeabilized with 0.1% Triton X-100 in PBS for another 15 minutes at room temperature. Cells were blocked with 4% horse serum in PBS for 1 hour and then incubated with primary antibodies overnight at 4 °C followed by incubation with appropriate fluorogenic secondary antibodies (Thermo Fisher Scientific) at room temperature for 1 hour.

### Quantitative real time PCR (qPCR)

Total RNAs from all cells were extracted using Trizol Reagent (Thermo Fisher Scientific) following the manufacturer’s instructions. RNA integrity was determined using formaldehyde denaturalization agarose gel electrophoresis. RNA concentrations were measured with Nanodrop spectrophotometer (Thermo Fisher Scientific). RNA was reverse transcribed by using iScript cDNA Synthesis Kit (BioRad). qPCR was performed using StepOne Real-Time PCR System (Thermo Fisher Scientific). Primer oligonucleotides were synthesized by Sigma and are listed in Supplementary Table S5.

### Spontaneous beating and calcium transient assessment

Spontaneous beating assessment was performed by light microscopy at room temperature after indicated treatment of MEFs or NCFs at serveral time points. Beating cell number was manually quantified by single-blind method in each well of 48-well plate. αMHC-Cre/Rosa26A-Flox-Stop-Flox-Gcamp3 NCFs were used for calcium transient assessment as previously reported (*41*) that was performed by fluorescence microscopy at room temperature after indicated reprogramming treatments. Three HPFs of view were randomly selected within each well.

### RNA-sequencing

Total RNAs from all group of cells were isolated using TRIzol following the provider’s instructions. RNA (RIN > 8.5) was used for RNA-seq library preparation by using NEBNext® Ultra™ II Directional RNA Library Prep Kit for Illumina (E7760S). The libraries were sequenced using Hiseq 4000 by the University of Michigan Sequencing Core. The quantification of RNA expression was estimated by Kallisto ^46^. The transcript abundance was imported using tximport followed by DESeq2 analysis. The abundance of genes was used to calculate fold change and p values. Differentially expressed genes were identified as a fold change greater than 2 and padj values less than 0.05.

### Condensation and optoIDR assay

The condensate regulation of IDRs of Hdac4, MEF2C and Foxo1 were predicted by dSCOPE, which is a web server developed for detecting sequences critical for phase separation related proteins (*61, 62*). Predicted IDRs on Hdac4 were deleted respectively, and each truncated Hdac4 constructs were inserted into mammalian expression vector pMX. Each pMX plasmids were transformed into HEK293T cells to test the condensate formation of each truncated Hdac4 construct.

The photoactivatable, self-associating Cry2 protein was labeled with mCherry and fused to the intrinsically disordered regions (IDR) Hdac4, MEF2C and Foxo1. The fusion of the IDR of Hdac4, MEF2C and Foxo1 to Cry2-mCherry were tested for the rapid formation of micron-sized spherical droplets upon blue light stimulation (*43*). WT and Mutated fusion protein were compared for their condensate formation ability by blue light stimulation within 10 minutes.

### Statistic analysis

Results were presented as mean ±s.e.m. Statistical difference between groups was analyzed by one-way ANOVA followed by the Student–Newman–Keuls multiple comparisons tests. A *P*-value <0.05 was regarded as significant.

---

**Fig. S1.**
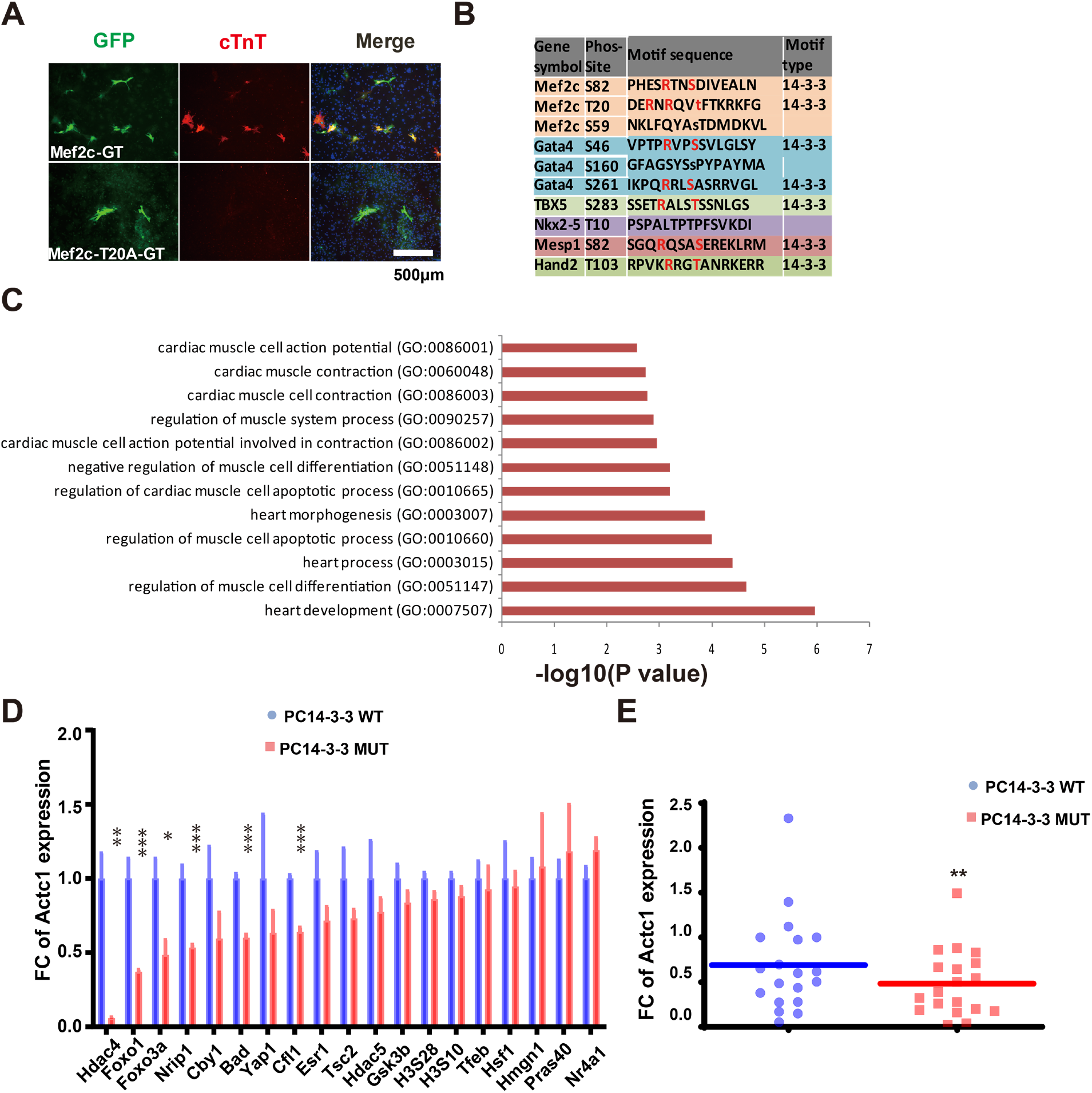
(A) Immunocytochemistry (ICC) of cardiac markers cTnt by fluorescence microscopy with over-expression of Mef2c T20A (D). (B) Listed results of the mutated site and cardiac TFs affect reprogramming. 14-3-3 binding motif was shown as the dominant motif in these TFs to regulate reprogramming. (C) Gene Ontology (GO) enrichment analysis of 14-3-3 binding proteins. Enriched pathways were listed based on P-value. (D) Relative expression of Actc1 in iCM reprogramming with WT and mutant PC14-3-3-containing proteins 14 days after MGT transduction. **P<0.01, vs WT. (E) Relative expression of Myh6 in iCM reprogramming with WT and mutant PC14-3-3-containing proteins 14 days after MGT transduction, Data are normalized to MGT+empty plasmids group. *P<0.05, **P<0.01, ***P<0.001, vs WT. FC : fold change

**Fig. S2.**
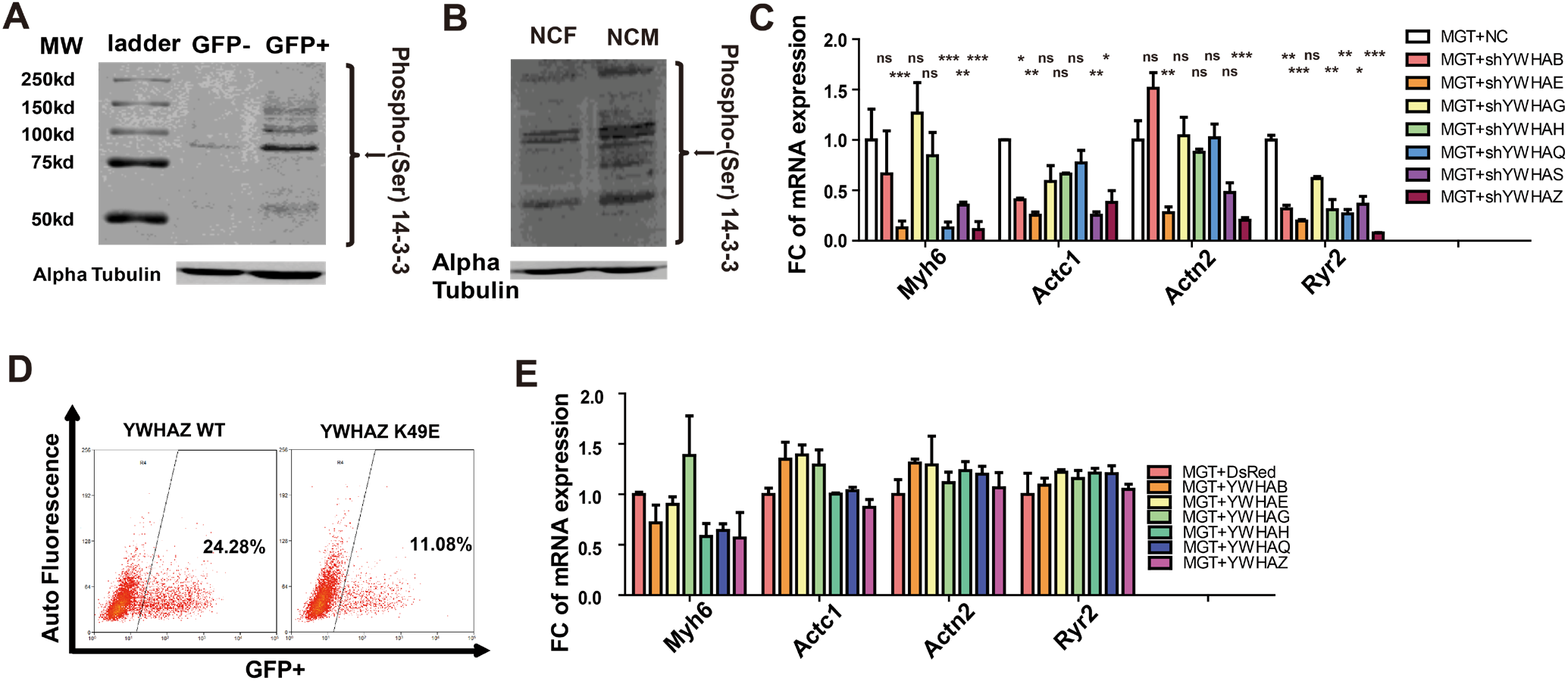
(A) Detection of 14-3-3 binding motif phosphorylation by Western blot of protein lysates from successful reprogrammed Myh6-GFP+ cells and the unsuccessful reprogrammed Myh6-GFP-cells separated by flow cytometry. (B) Detection of 14-3-3 binding motif phosphorylation by Western blot of protein lysates from neonatal cardiomyocyte and neonatal cardiac fibroblast. *P<0.05, **P<0.01, ***P<0.001, vs NC. (C) The mRNA change of direct shRNA knockdown of 7 14-3-3 isoforms to MGT infection MEF cells by QPCR. *P<0.05, **P<0.01, ***P<0.001, vs NC. (D) Flow cytometry analysis of Myh6-GFP+ cells in inducible cardiac reprogramming cell line by expression WT or K49E mutation YWHAE in the reprogramming process (E) Relative expression of cardiomyocyte (CM) marker genes in iCM reprogramming under the treatment of 14-3-3 isoforms overexpression after MGT transduction.

**Fig. S3.**
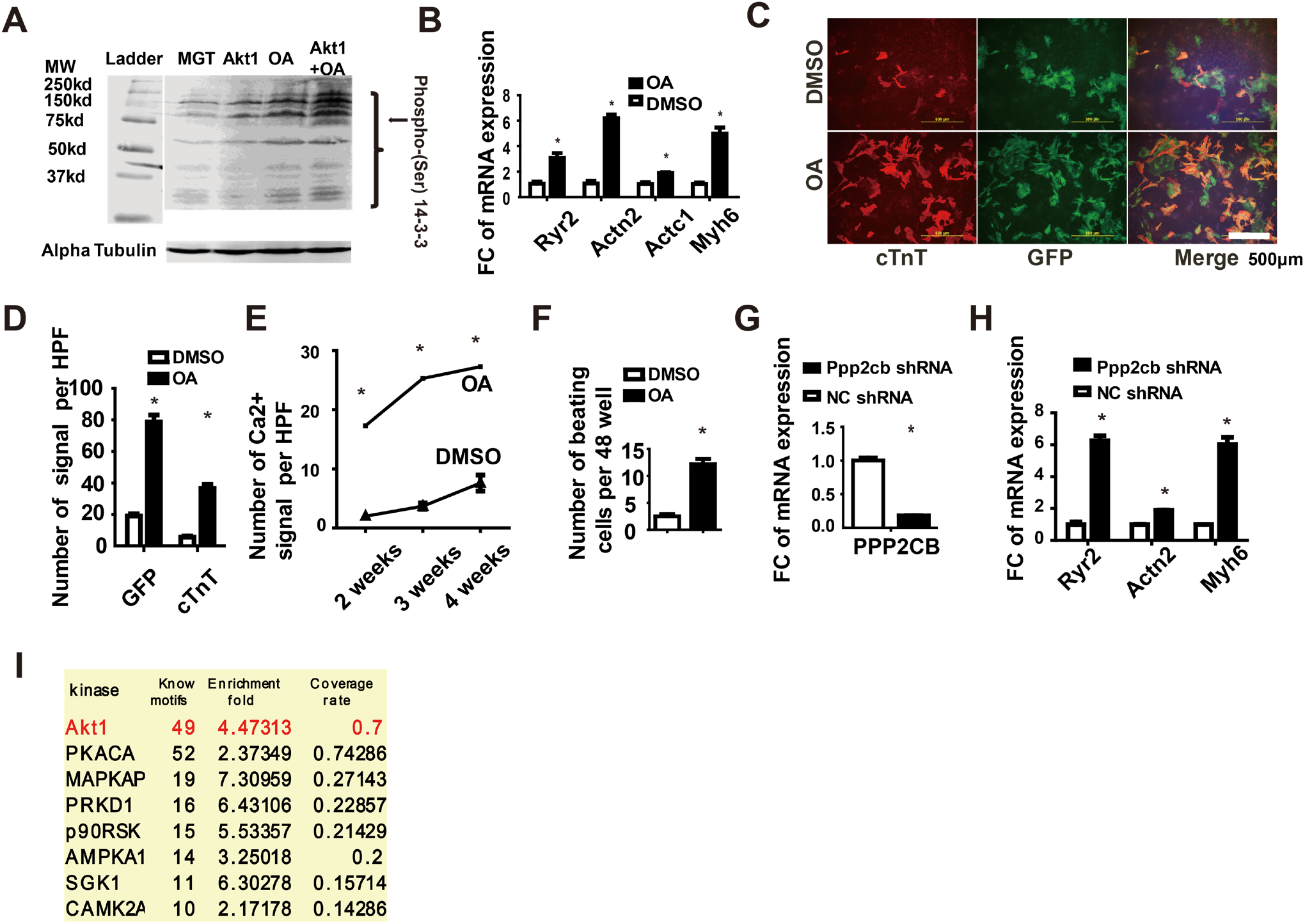
(A) Combined effect of Akt1 and OA treatment on 14-3-3 binding motif phosphorylation. Phosphorylation of 14-3-3 binding motifs was detected by Western blot of protein lysates from MEF cells with indicated treatments. (B) Relative expression of cardiomyocyte (CM) marker genes of iCM reprogramming with OA treatment 14 days after MGT transduction. *P<0.05, **P<0.01, ***P<0.001, vs DMSO. (C&D) Immunocytochemistry (ICC) and Quantification of cardiac markers Tnt and Myh6-GFP of MGT-transduced cells with or without OA treatment by fluorescence microscopy (100×). *P<0.05, **P<0.01, ***P<0.001, vs DMSO. (E) Quantification of spontaneous Ca^2+^ oscillations cells per field with or without OA treatment for 1 to 4 weeks (n=50 from 10 wells). *P<0.05, **P<0.01, ***P<0.001, vs DMSO. (F) Quantification of beating iCMs loci with indicated viral infection for 4 weeks (n=10). *P<0.05, **P<0.01, ***P<0.001, vs DMSO. (G) Relative expression of Ppp2cb in iCM reprogramming with shRNA treatment. *P<0.05, **P<0.01, ***P<0.001, vs NC. (H) Relative expression of cardiomyocyte (CM) marker genes in iCM reprogramming with or without shRNA Ppp2cb treatment 14 days treatment after MGT transduction. *P<0.05, **P<0.01, ***P<0.001, vs NC. (I) Candidate kinases that phosphorylate 14-3-3 binding motifs during reprogramming and the coverage of those kinase for 14-3-3 binding motif in different protein.

**Fig. S4.**
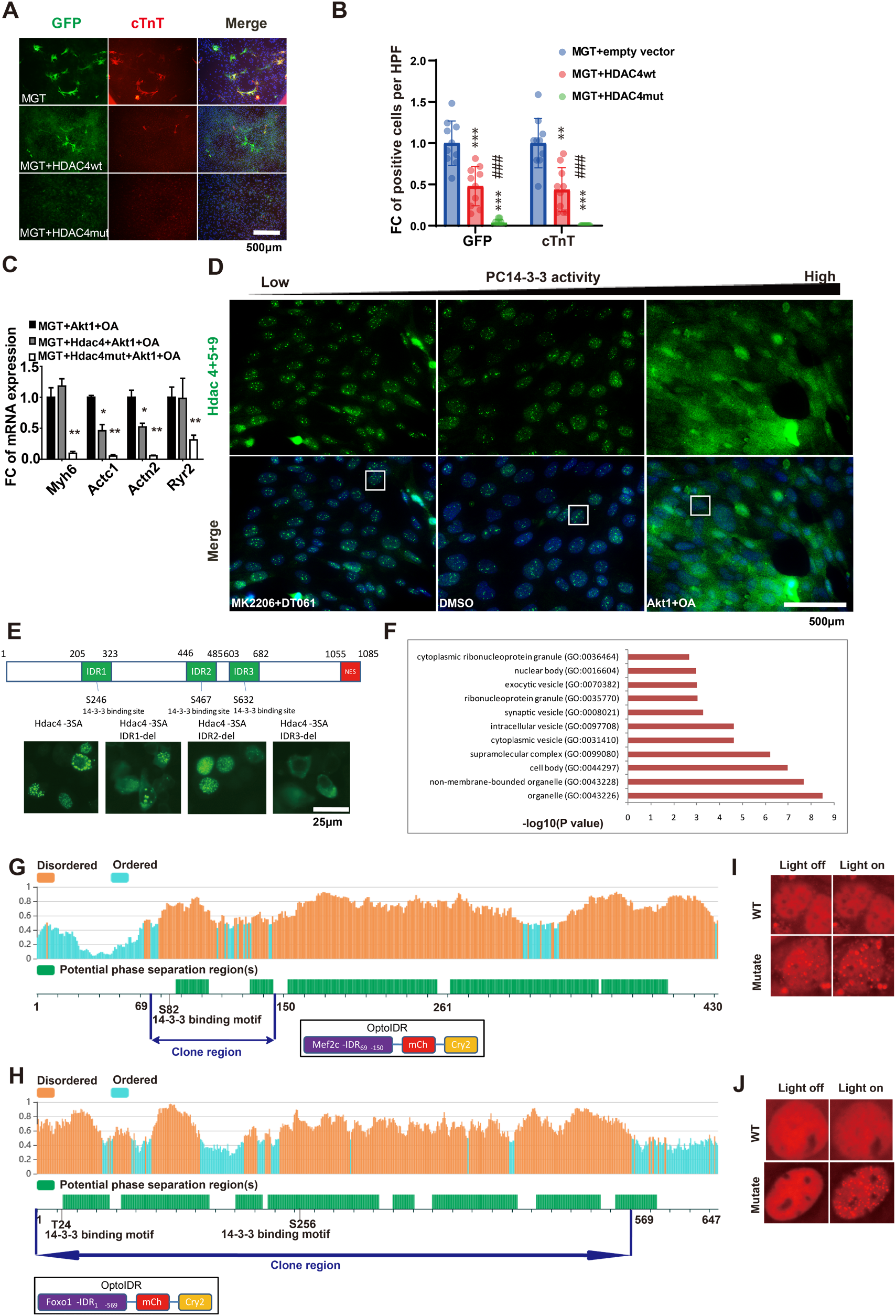
(A&B) Immunocytochemistry (ICC) and quantification of cardiac markers Myh6-GFP and Tnnt2 of Hdac4 wt or mut-transduced cells by fluorescence microscopy. FC : fold change. (C) Relative expression of cardiomyocyte marker genes in iCM reprogramming with WT and mutant Hdac4 14 days after MGT transduction with Akt1 and OA treatment. (D) Immunocytochemistry of endogenous Hdac4 in reprogramming cells treated with OA + Akt1 or MK-2206 (Akt1 inhibitor)+ DT-061 (PP2A activator) by fluorescence microscopy. (E) A schematic diagram of Hdac4 disordered region deletion strategy, the 14-3-3 binding motif and adjacent IDR region on Hdac4 was deleted, the GFP-Hdac4-Flag living cell condensates were observered, deleted 14-3-3 binding motif and adjacent region IDR cause lost Hdac4 condensate formation. (F) Pathway analysis or Gene Ontology (GO) enrichment analysis to identify phosphorylation of 14-3-3 binding motif proteins. The proteins related with pathways related with nuclear bodies were listed. (G-H) A schematic diagram of MEF2C (E) and Foxo1 (F) disordered region, potential phase separation regions and the construction strategy for optoIDR assay. (I-J) Images of the HE293T cell expressing MEF2C or Foxo1-optoIDR subjected to laser excitation for the times indicated. A droplet fusion event occurs in the MEF2Cmut or Foxo1mut-optoIDR optoIDR group but not MEF2Cwt or Foxo1wt-optoIDR.

**Fig. S5.**
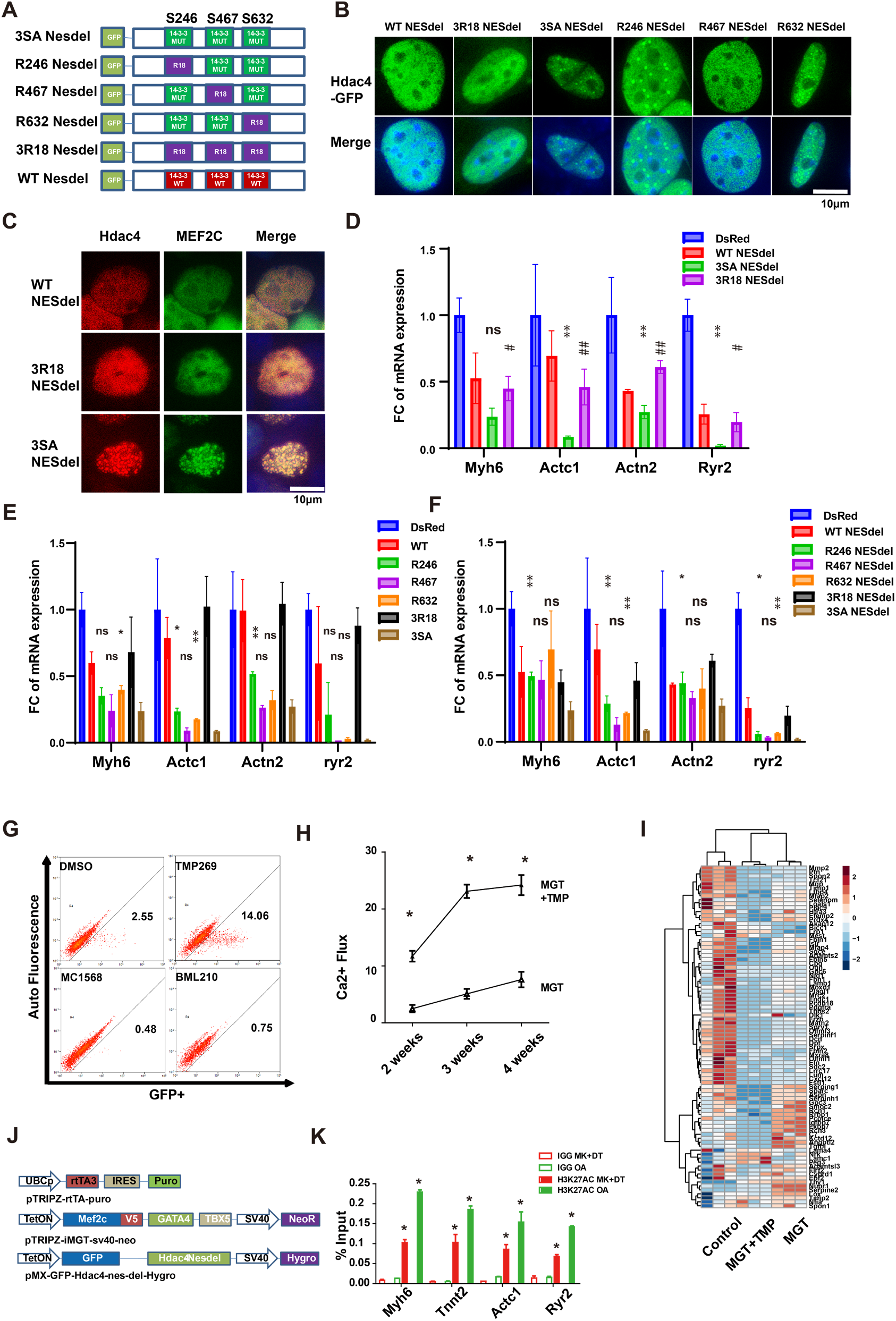
(A) A schematic diagram of 14-3-3 permanent bind form of Hdac4 and 14-3-3 binding motif mutation Hdac4 and WT without NES. (B) Images of the reprogramming cell expression of 14-3-3 permanent bind form of Hdac4 and control without NES. (C) Images of the MEF cell expression of 14-3-3 permanent bind (R18) and control forms of Hdac4 without NES with MGT. (D) Relative expression of cardiomyocyte (CM) marker genes of iCM reprogramming with expression of 14-3-3 permanent bind form of Hdac4 without NES. *P<0.05, **P<0.01, ***P<0.001, WT vs 3SA. #P<0.05, ##P<0.01, ###P<0.001, 3SA vs 3R18. (E) Relative expression of cardiomyocyte (CM) marker genes of iCM reprogramming with expression of 14-3-3 permanent bind form of Hdac4 with single R18 insertion. *P<0.05, **P<0.01, ***P<0.001, vs 3SA. (F) Relative expression of cardiomyocyte (CM) marker genes of iCM reprogramming with expression of 14-3-3 permanent bind form of Hdac4 with single R18 insertion without NES. *P<0.05, **P<0.01, ***P<0.001, vs 3SA. (G) Successful reprogrammed Myh6-GFP+ cells was detected by flow cytometry treatment with t Hdac4 inhibitors, TMP269, MC1568 and BML210. (H) Quantification of spontaneous Ca^2+^ oscillations cells per field with TMP269 treatment for 2 to 4 weeks (n=50 from 10 wells). *P<0.05, **P<0.01, ***P<0.001. (I) Heat map showing differential expression of representative fibroblast-related genes among MGT-transduced cells with or without Hdac4 inhibitor TMP269 treatment. (J) A schematic diagram for the establishment of an inducible cardiac reprogramming cell line. (K) ChIP-qPCR for H3K27ac at cardiac gene loci of iMGT cells, treated with PC14-3-3 activation OA treatment or PC14-3-3 repressive treatment MK2206+DT-061.

**Table S1.**
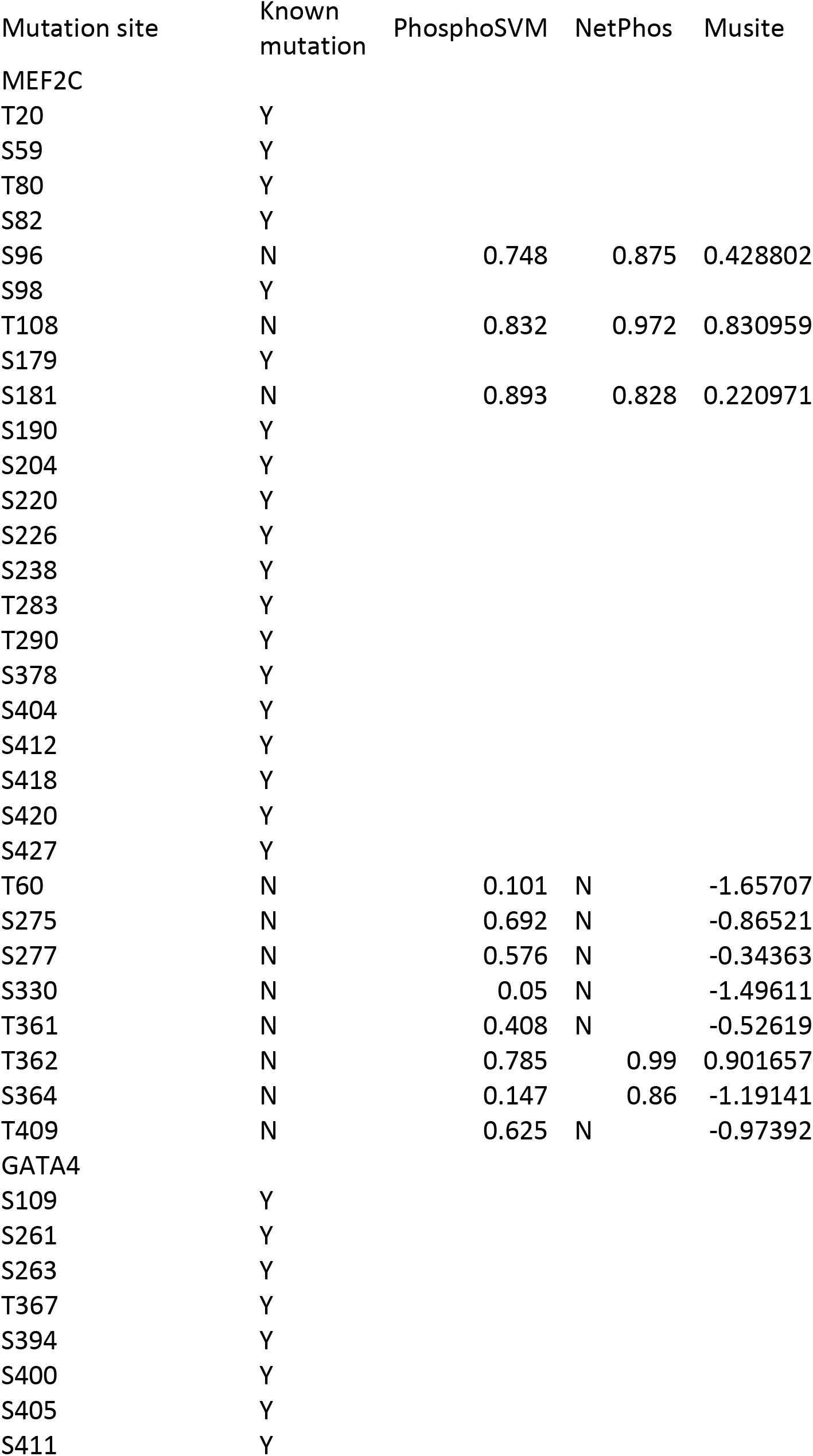
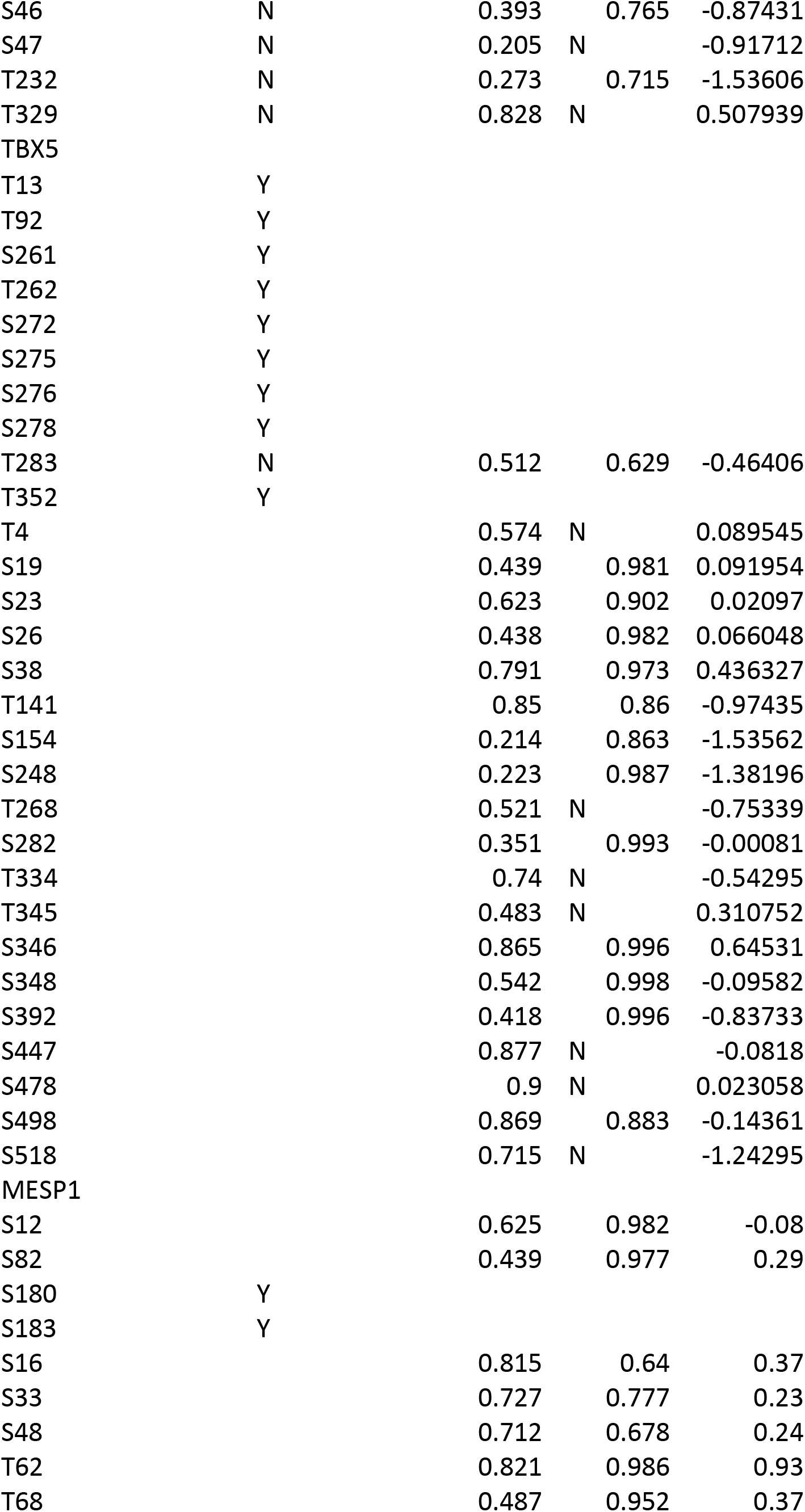
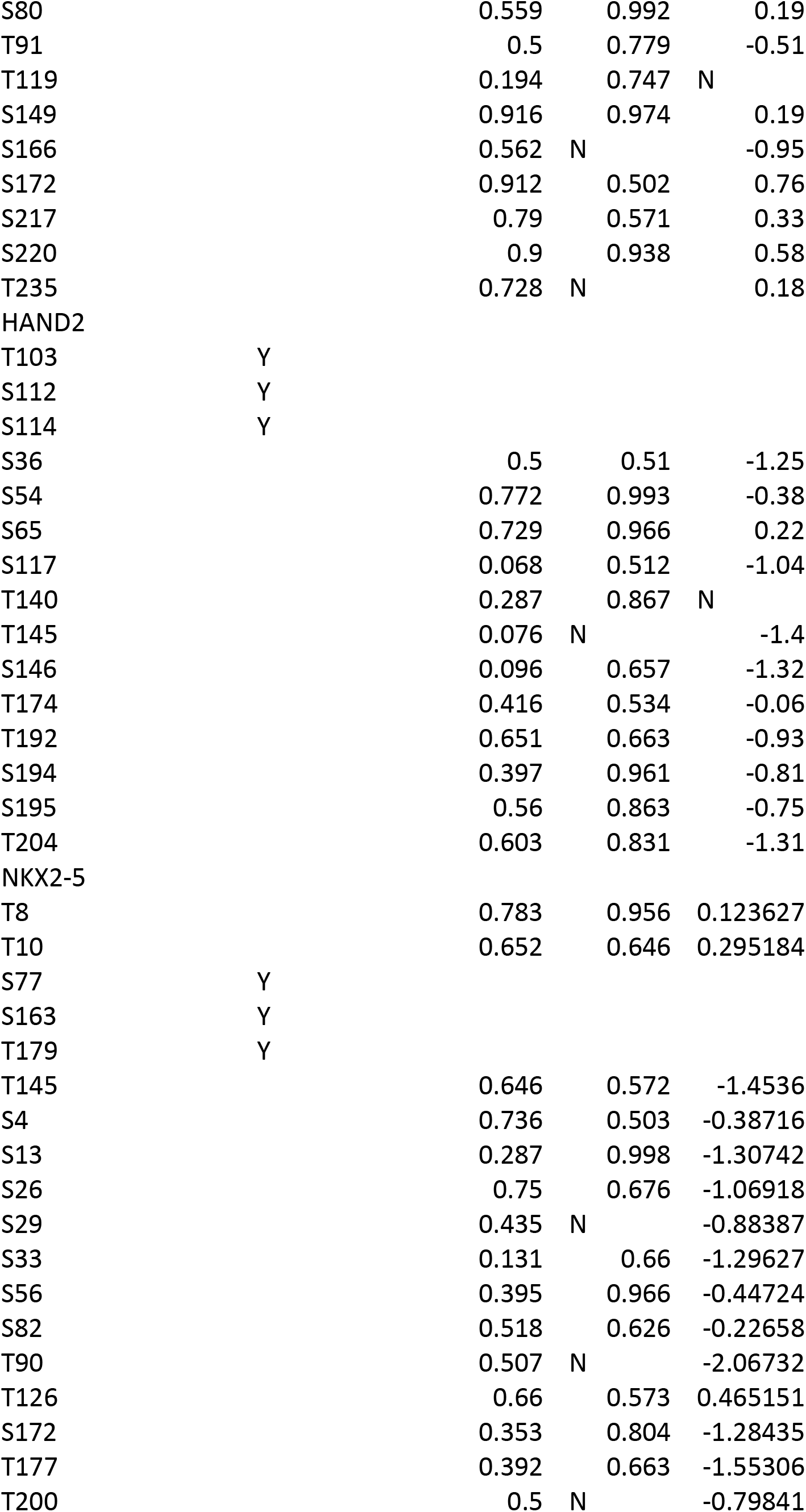
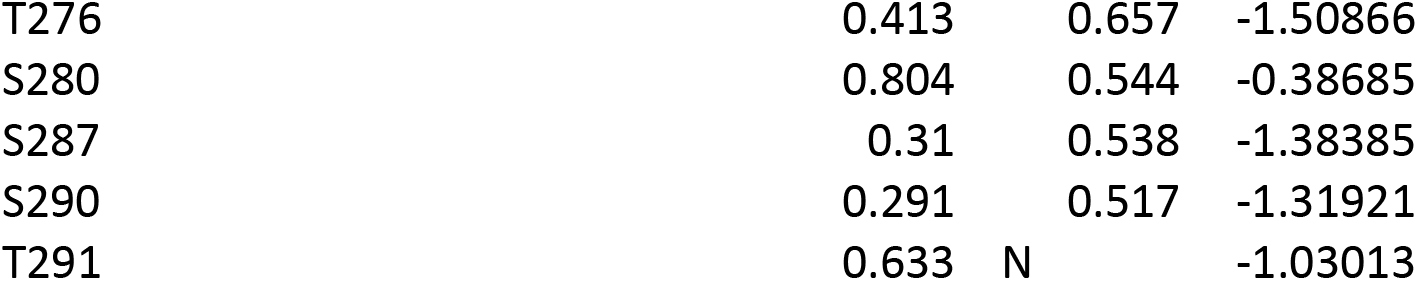
Predicted phosphorylation sites by PhosphoSVM, NetPhos and Musite.

| Mutation site | Known mutation | PhosphoSVM | NetPhos | Musite |
| --- | --- | --- | --- | --- |
| MEF2C |  |  |  |  |
| T20 | Y |  |  |  |
| S59 | Y |  |  |  |
| T80 | Y |  |  |  |
| S82 | Y |  |  |  |
| S96 | N | 0.748 | 0.875 | 0.428802 |
| S98 | Y |  |  |  |
| T108 | N | 0.832 | 0.972 | 0.830959 |
| S179 | Y |  |  |  |
| S181 | N | 0.893 | 0.828 | 0.220971 |
| S190 | Y |  |  |  |
| S204 | Y |  |  |  |
| S220 | Y |  |  |  |
| S226 | Y |  |  |  |
| S238 | Y |  |  |  |
| T283 | Y |  |  |  |
| T290 | Y |  |  |  |
| S378 | Y |  |  |  |
| S404 | Y |  |  |  |
| S412 | Y |  |  |  |
| S418 | Y |  |  |  |
| S420 | Y |  |  |  |
| S427 | Y |  |  |  |
| T60 | N | 0.101 | N | -1.65707 |
| S275 | N | 0.692 | N | -0.86521 |
| S277 | N | 0.576 | N | -0.34363 |
| S330 | N | 0.05 | N | -1.49611 |
| T361 | N | 0.408 | N | -0.52619 |
| T362 | N | 0.785 | 0.99 | 0.901657 |
| S364 | N | 0.147 | 0.86 | -1.19141 |
| T409 | N | 0.625 | N | -0.97392 |
| GATA4 |  |  |  |  |
| S109 | Y |  |  |  |
| S261 | Y |  |  |  |
| S263 | Y |  |  |  |
| T367 | Y |  |  |  |
| S394 | Y |  |  |  |
| S400 | Y |  |  |  |
| S405 | Y |  |  |  |
| S411 | Y |  |  |  |
| S46 | N | 0.393 | 0.765 | -0.87431 |
| S47 | N | 0.205 | N | -0.91712 |
| T232 | N | 0.273 | 0.715 | -1.53606 |
| T329 | N | 0.828 | N | 0.507939 |
| TBX5 |  |  |  |  |
| T13 | Y |  |  |  |
| T92 | Y |  |  |  |
| S261 | Y |  |  |  |
| T262 | Y |  |  |  |
| S272 | Y |  |  |  |
| S275 | Y |  |  |  |
| S276 | Y |  |  |  |
| S278 | Y |  |  |  |
| T283 | N | 0.512 | 0.629 | -0.46406 |
| T352 | Y |  |  |  |
| T4 |  | 0.574 | N | 0.089545 |
| S19 |  | 0.439 | 0.981 | 0.091954 |
| S23 |  | 0.623 | 0.902 | 0.02097 |
| S26 |  | 0.438 | 0.982 | 0.066048 |
| S38 |  | 0.791 | 0.973 | 0.436327 |
| T141 |  | 0.85 | 0.86 | -0.97435 |
| S154 |  | 0.214 | 0.863 | -1.53562 |
| S248 |  | 0.223 | 0.987 | -1.38196 |
| T268 |  | 0.521 | N | -0.75339 |
| S282 |  | 0.351 | 0.993 | -0.00081 |
| T334 |  | 0.74 | N | -0.54295 |
| T345 |  | 0.483 | N | 0.310752 |
| S346 |  | 0.865 | 0.996 | 0.64531 |
| S348 |  | 0.542 | 0.998 | -0.09582 |
| S392 |  | 0.418 | 0.996 | -0.83733 |
| S447 |  | 0.877 | N | -0.0818 |
| S478 |  | 0.9 | N | 0.023058 |
| S498 |  | 0.869 | 0.883 | -0.14361 |
| S518 |  | 0.715 | N | -1.24295 |
| MESP1 |  |  |  |  |
| S12 |  | 0.625 | 0.982 | -0.08 |
| S82 |  | 0.439 | 0.977 | 0.29 |
| S180 | Y |  |  |  |
| S183 | Y |  |  |  |
| S16 |  | 0.815 | 0.64 | 0.37 |
| S33 |  | 0.727 | 0.777 | 0.23 |
| S48 |  | 0.712 | 0.678 | 0.24 |
| T62 |  | 0.821 | 0.986 | 0.93 |
| T68 |  | 0.487 | 0.952 | 0.37 |
| S80 |  | 0.559 | 0.992 | 0.19 |
| T91 |  | 0.5 | 0.779 | -0.51 |
| T119 |  | 0.194 | 0.747 | N |
| S149 |  | 0.916 | 0.974 | 0.19 |
| S166 |  | 0.562 | N | -0.95 |
| S172 |  | 0.912 | 0.502 | 0.76 |
| S217 |  | 0.79 | 0.571 | 0.33 |
| S220 |  | 0.9 | 0.938 | 0.58 |
| T235 |  | 0.728 | N | 0.18 |
| HAND2 |  |  |  |  |
| T103 | Y |  |  |  |
| S112 | Y |  |  |  |
| S114 | Y |  |  |  |
| S36 |  | 0.5 | 0.51 | -1.25 |
| S54 |  | 0.772 | 0.993 | -0.38 |
| S65 |  | 0.729 | 0.966 | 0.22 |
| S117 |  | 0.068 | 0.512 | -1.04 |
| T140 |  | 0.287 | 0.867 | N |
| T145 |  | 0.076 | N | -1.4 |
| S146 |  | 0.096 | 0.657 | -1.32 |
| T174 |  | 0.416 | 0.534 | -0.06 |
| T192 |  | 0.651 | 0.663 | -0.93 |
| S194 |  | 0.397 | 0.961 | -0.81 |
| S195 |  | 0.56 | 0.863 | -0.75 |
| T204 |  | 0.603 | 0.831 | -1.31 |
| NKX2-5 |  |  |  |  |
| T8 |  | 0.783 | 0.956 | 0.123627 |
| T10 |  | 0.652 | 0.646 | 0.295184 |
| S77 | Y |  |  |  |
| S163 | Y |  |  |  |
| T179 | Y |  |  |  |
| T145 |  | 0.646 | 0.572 | -1.4536 |
| S4 |  | 0.736 | 0.503 | -0.38716 |
| S13 |  | 0.287 | 0.998 | -1.30742 |
| S26 |  | 0.75 | 0.676 | -1.06918 |
| S29 |  | 0.435 | N | -0.88387 |
| S33 |  | 0.131 | 0.66 | -1.29627 |
| S56 |  | 0.395 | 0.966 | -0.44724 |
| S82 |  | 0.518 | 0.626 | -0.22658 |
| T90 |  | 0.507 | N | -2.06732 |
| T126 |  | 0.66 | 0.573 | 0.465151 |
| S172 |  | 0.353 | 0.804 | -1.28435 |
| T177 |  | 0.392 | 0.663 | -1.55306 |
| T200 |  | 0.5 | N | -0.79841 |

|  |  |  |  |
| --- | --- | --- | --- |
| T276 | 0.413 | 0.657 | -1.50866 |
| S280 | 0.804 | 0.544 | -0.38685 |
| S287 | 0.31 | 0.538 | -1.38385 |
| S290 | 0.291 | 0.517 | -1.31921 |
| T291 | 0.633 | N | -1.03013 |

**Table S2.**
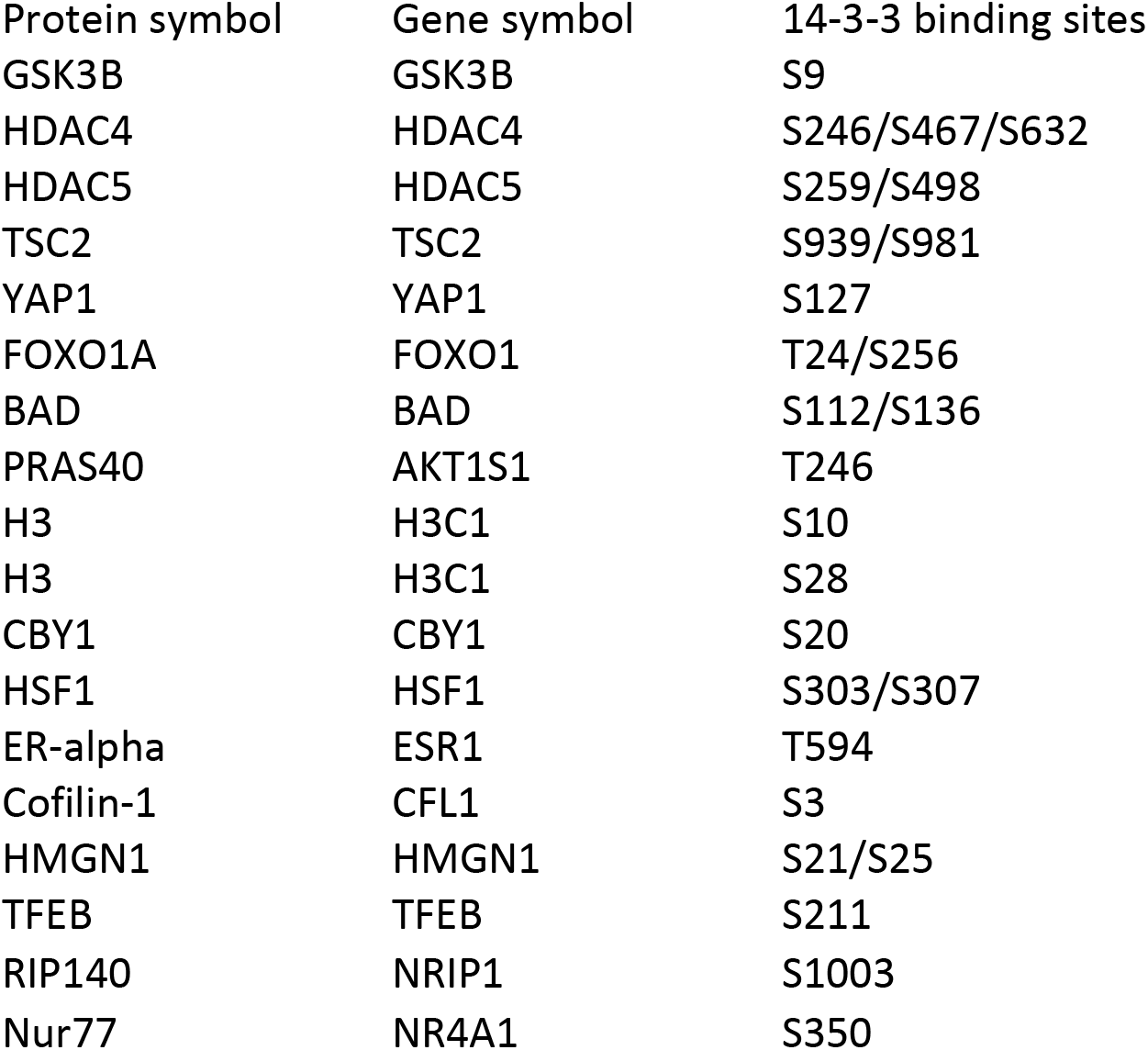
19 key transcription factors, epigenetic factors and histone proteins carrying known 14-3-3 binding motifs.

**Table S3.**
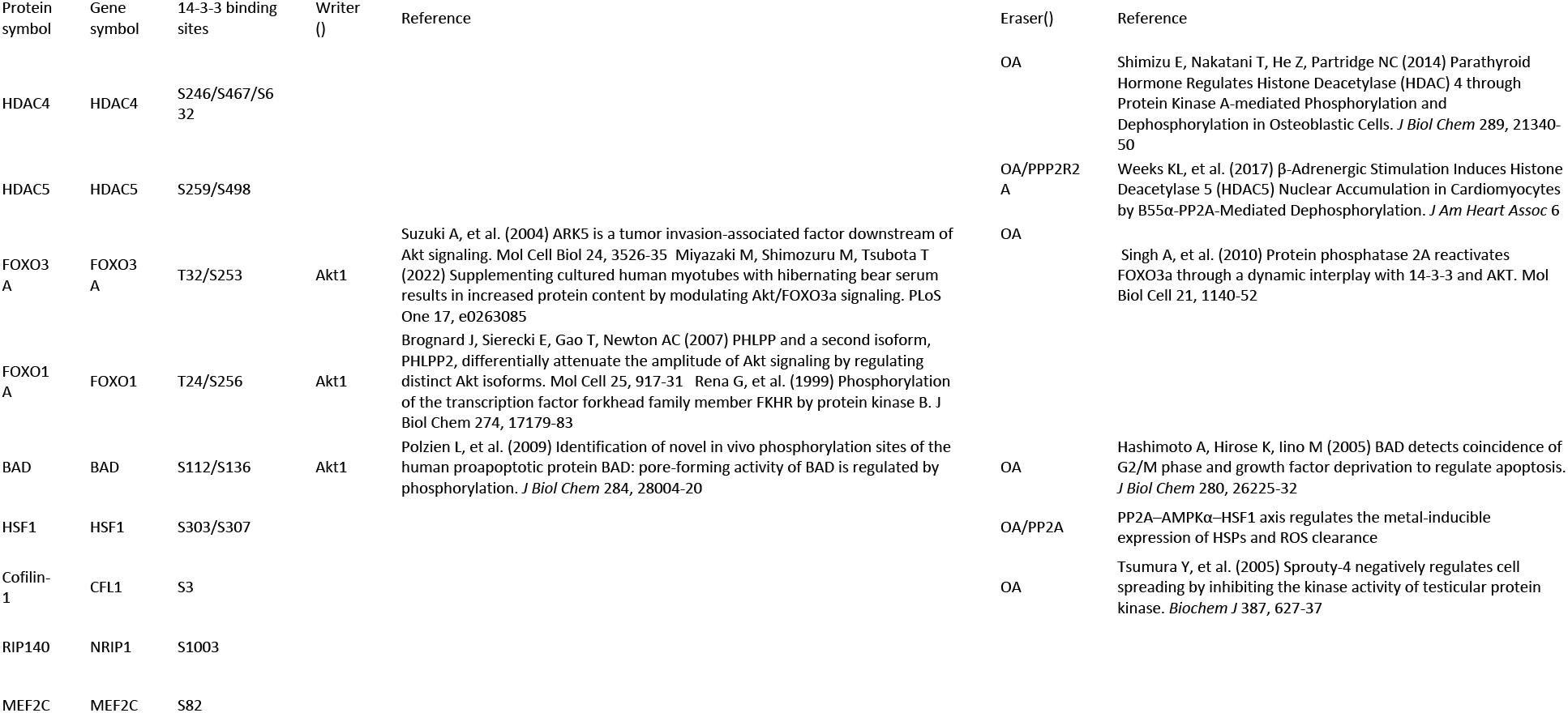
14-3-3 binding motif embedded proteins identified in Fig. 1G that affected reprogramming regulated by Akt1 or PP2A.

**Table S4.**
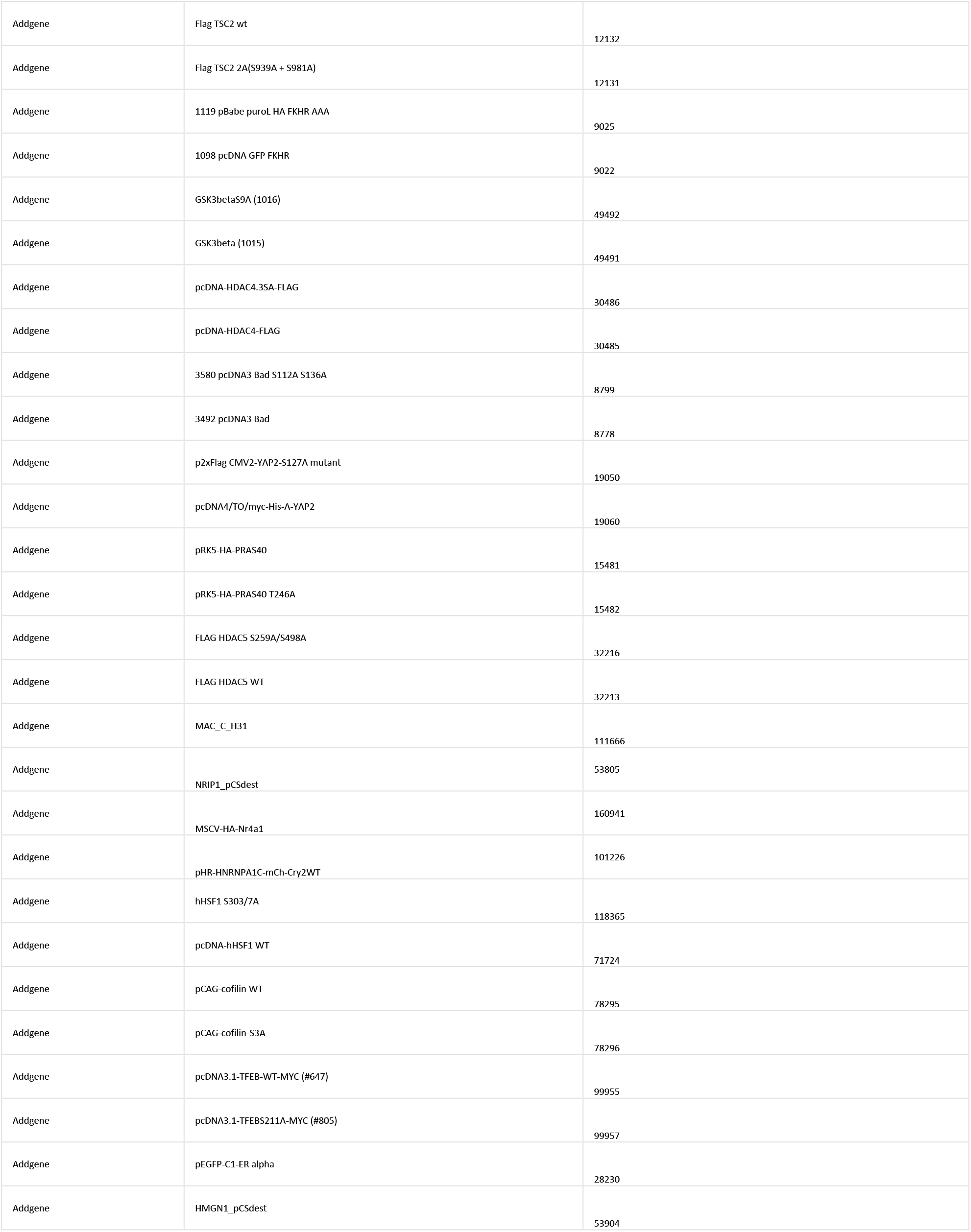
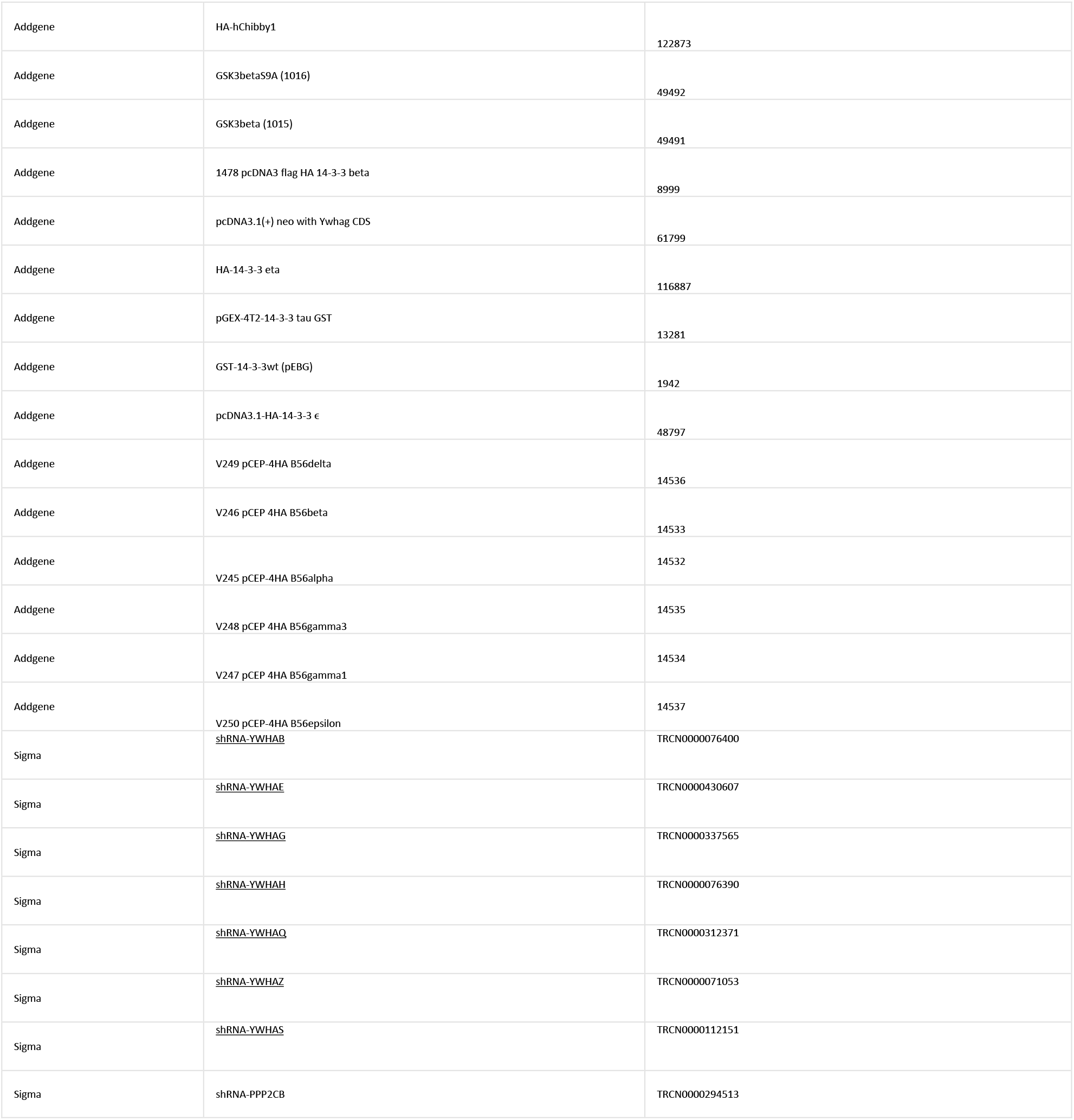
The plasmids purchase from Addgene and Sigma.

**Table S5.**
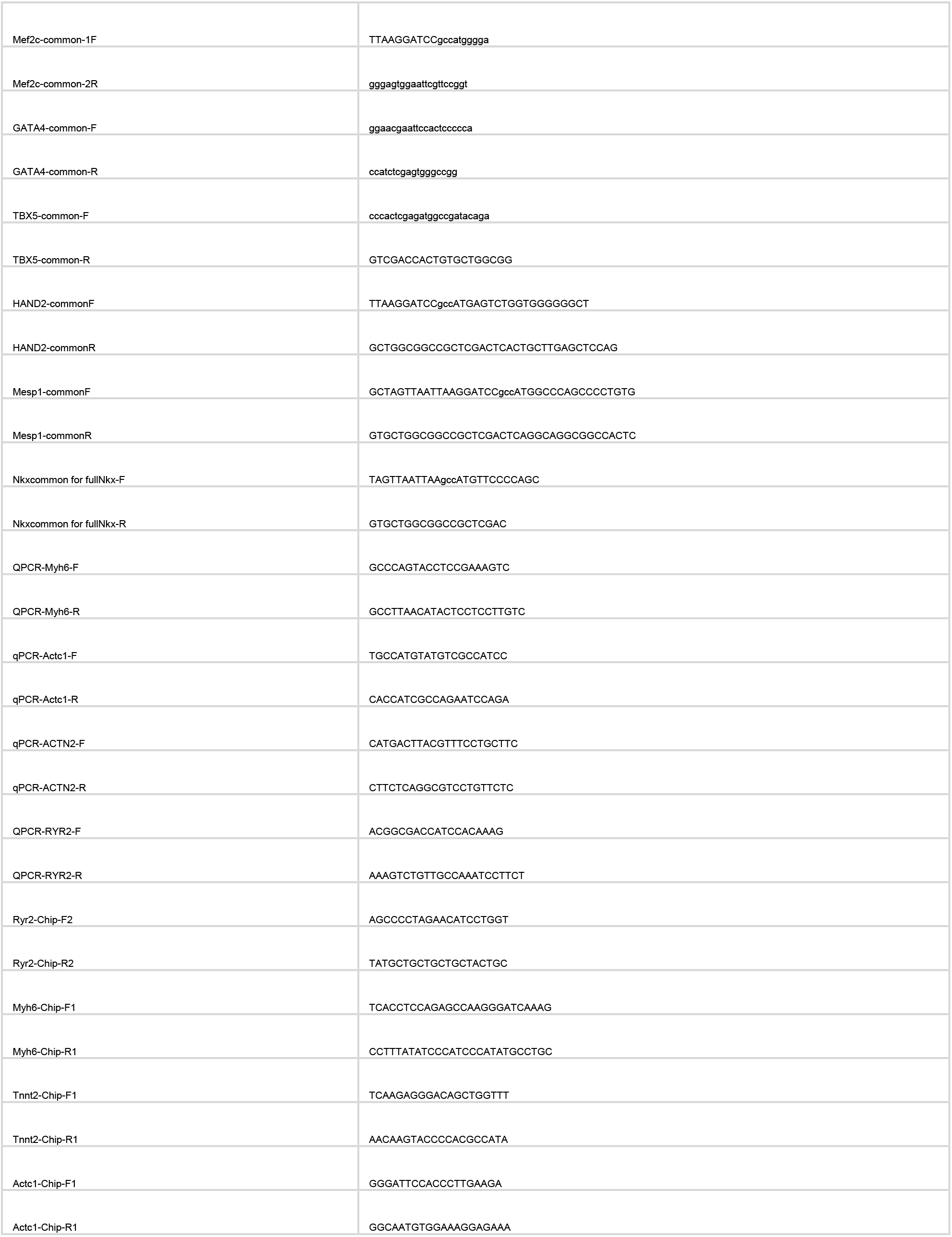
The main primers used for the cloning.

## Notes

### Competing Interest Statement

The authors have declared no competing interest.

